# Automatic design of gene regulatory mechanisms for spatial pattern formation

**DOI:** 10.1101/2023.07.26.550573

**Authors:** Reza Mousavi, Daniel Lobo

**Affiliations:** Department of Biological Sciences, University of Maryland, Baltimore County, 1000 Hilltop Circle, Baltimore, MD 21250, USA; Greenebaum Comprehensive Cancer Center and Center for Stem Cell Biology & Regenerative Medicine, University of Maryland, School of Medicine, 22 S. Greene Street, Baltimore, MD 21201, USA

**Keywords:** Gene Expression Patterns, Gene Regulatory Mechanisms, Systems Biology, Machine Learning, Synthetic Biology, Evolutionary Computation

## Abstract

Synthetic developmental biology aims to engineer gene regulatory mechanisms (GRMs) for understanding and producing desired multicellular patterns and shapes. However, designing GRMs for spatial patterns is a current challenge due to the nonlinear interactions and feedback loops in genetic circuits. Here we present a methodology to automatically design GRMs that can produce any given spatial pattern. The proposed approach uses two orthogonal morphogen gradients acting as positional information signals in a multicellular tissue area or culture, which constitutes a continuous field of engineered cells implementing the same designed GRM. To efficiently design both the circuit network and the interaction mechanisms—including the number of genes necessary for the formation of the target pattern—we developed an automated algorithm based on high-performance evolutionary computation. The tolerance of the algorithm can be configured to design GRMs that are either simple to produce approximate patterns or complex to produce precise patterns. We demonstrate the approach by automatically designing GRMs that can produce a diverse set of synthetic spatial expression patterns by interpreting just two orthogonal morphogen gradients. The proposed framework offers a versatile approach to systematically design and discover pattern-producing genetic circuits.

## 1. Introduction

Gene regulatory mechanisms (GRMs) comprise a network of genes and signal molecules together with their mechanistic interactions. They can govern the development of gene expression patterns in time and space, which in turn can control the formation of anatomical structures, organ locations, and complex shapes (Kicheva and Briscoe, 2015). Understanding the regulatory mechanisms that can produce particular spatial patterns is crucial for the prediction of phenotypes and the identification of interventions toward desired outputs (Lobo et al., 2017). Furthermore, the design and engineering of complex GRMs in synthetic developmental biology (Santos-Moreno and Schaerli, 2019; Zarkesh et al., 2022) will allow the manufacturing of multicellular patterns, value-added bioproducts, and synthetic smart biomaterials for medical and industrial applications (Barbier et al., 2022; Basu et al., 2005; Kim et al., 2020; Liu et al., 2011). Towards this, automated algorithms have been proposed to aid the molecular engineering of pre-designed synthetic gene circuits (Appleton et al., 2017; Buecherl and Myers, 2022; Nielsen et al., 2016) as well as for the design of novel circuits that can implement a given simple behavior, such as switches, oscillators, or basic functions (Dasika and Maranas, 2008; Hiscock, 2019; Huynh et al., 2013; Marchisio and Stelling, 2011; Rodrigo and Jaramillo, 2013; Rodrigo et al., 2011). However, automatically discovering or designing a GRM that can produce complex behaviors, such as a target spatial expression pattern, is still a current major challenge (Ko et al., 2022; Stillman and Mayor, 2023). Although random search has been used for exploring and engineering GRMs for simple spatial patterns such as a stripe (Cotterell and Sharpe, 2010; Schaerli et al., 2014), heuristic optimization methods are required for automatically designing complex GRMs (Delgado and Gómez-Vela, 2019).

Different heuristic methods exist for reconstructing complex gene regulatory networks (GRNs)— a network of predicted gene-gene regulatory links lacking mechanistic information. These approaches infer links in the network as probabilistic gene-gene interactions (Zhou and Cai, 2020), typically from large-scale transcriptomics data. Unsupervised machine learning can take unlabeled data from experimental or synthetic nonspatial gene expression patterns, such as microarray or RNA-Seq transcriptomics, to predict GRNs (Aibar et al., 2017; Faith et al., 2007; Haury et al., 2012; Huynh-Thu et al., 2010; Margolin et al., 2006; Meyer et al., 2007; Petralia et al., 2015; Yin et al., 2006). In contrast, supervised machine learning uses known gene-gene interactions as labeled data for training the models and then infer GRNs from non-spatial transcriptomics data (Gillani et al., 2014; Maetschke et al., 2014; Mordelet and Vert, 2008; Razaghi-Moghadam and Nikoloski, 2020). In addition, automated methods have been proposed to infer GRNs from spatial gene expression patterns in *in situ* hybridization images, such as those during early *Drosophila* development (Wu et al., 2016; Yang et al., 2019). However, these GRN-inference methods are limited to inferring the structure (topology) of the gene regulatory network and hence lack the mechanistic details in the gene interactions essential for predicting spatial pattern formation dynamics. Indeed, inferring such GRMs needs computational methods based on dynamic mathematical formalisms that can mechanistically model signal- and gene-gene interactions to predict the resulting spatial phenotypes (Durant et al., 2016; Eskandari and Kuhl, 2015; Marcon and Sharpe, 2012).

Several heuristic optimization methods have been proposed for the inference of dynamic GRMs from spatial expression patterns (Ko et al., 2022). These pattern-forming GRMs are typically formalized with partial differential equations (PDEs) due to their ability to combine controlling mechanisms of gene regulation with spatial signaling and their resulting cellular behaviors in time and space (Cotterell and Sharpe, 2010; Schaerli et al., 2014). Evolutionary computation is a heuristic population-based method that can optimize complex solutions (Holland, 1975; Mousavi and Eftekhari, 2015; Mousavi et al., 2015; Mousavi et al., 2018; Reali et al., 2017). This evolutionary approach can infer the parameters of one-dimensional PDE models of development, such as for early Drosophila development (Jaeger et al., 2004a; Jaeger et al., 2004b; Verd et al., 2017), as well as the parameters and the structure of GRMs for one-dimensional embryonic dynamic patterns (Francois and Siggia, 2010; Henry et al., 2018) and simple two-dimensional planarian head-trunk-tail body patterns (Lobo and Levin, 2015; Lobo and Levin, 2017; Lobo et al., 2016). In addition, evolutionary computation has been proposed for designing synthetic gene circuits (Noman et al., 2013) that can perform basic tasks including switches (Francois and Hakim, 2004), logic gates (Rodrigo et al., 2007), oscillators (Smith et al., 2017), and a stripe pattern (Otero-Muras and Banga, 2017). However, the automatic design of GRMs that can produce an arbitrarily complex spatial pattern is still a current challenge.

Here, we present a novel methodology for the automatic design of GRMs that can dynamically produce a given spatial gene expression pattern in response to positional information gradient signals. The method leverages the advantages of evolutionary computation and high-performance computing (Mousavi et al., 2021) to rapidly design spatiotemporal GRMs. We defined a versatile set of non-linear gene regulatory mechanisms that serve as building blocks for the optimization method to design GRMs that develop the target pattern. Furthermore, the method can be tuned to produce either complex GRMs that develop precise patterns or simple GRMs that develop approximate patterns. We evaluate the performance of the methodology by successfully inferring GRMs for a diverse set of synthetic two-dimensional spatial patterns, including geometric shapes, symbols, and characters.

## 2. Results

### 2.1. A system for spatial pattern formation based on orthogonal gradient signals

Patterns in developmental biology are often formed by GRMs that react to diffusible morphogen signals producing spatial gradients (Stapornwongkul and Vincent, 2021). These signals act as a positional information system for cells to react differentially depending on their location (Tkačik and Gregor, 2021), a process that can be applied to synthetic spatial behaviors (Grant et al., 2016) and reinforced with synthetic bistable switches (Barbier et al., 2020). Here we employed a similar *in silico* approach based on continuous orthogonal morphogen gradients that serve as input signals for an automatically designed GRM to form a target spatial pattern (Fig. 1). The two input signals (labeled red and green) are produced from the top and left sides, respectively, of a two-dimensional cell culture domain and form similar but orthogonal static gradients. Each cell in the domain encodes the same GRM, which takes as input the two input signals and through a cascade of regulatory interactions expresses a non-diffusible reporter gene (blue).

**Figure 1.**
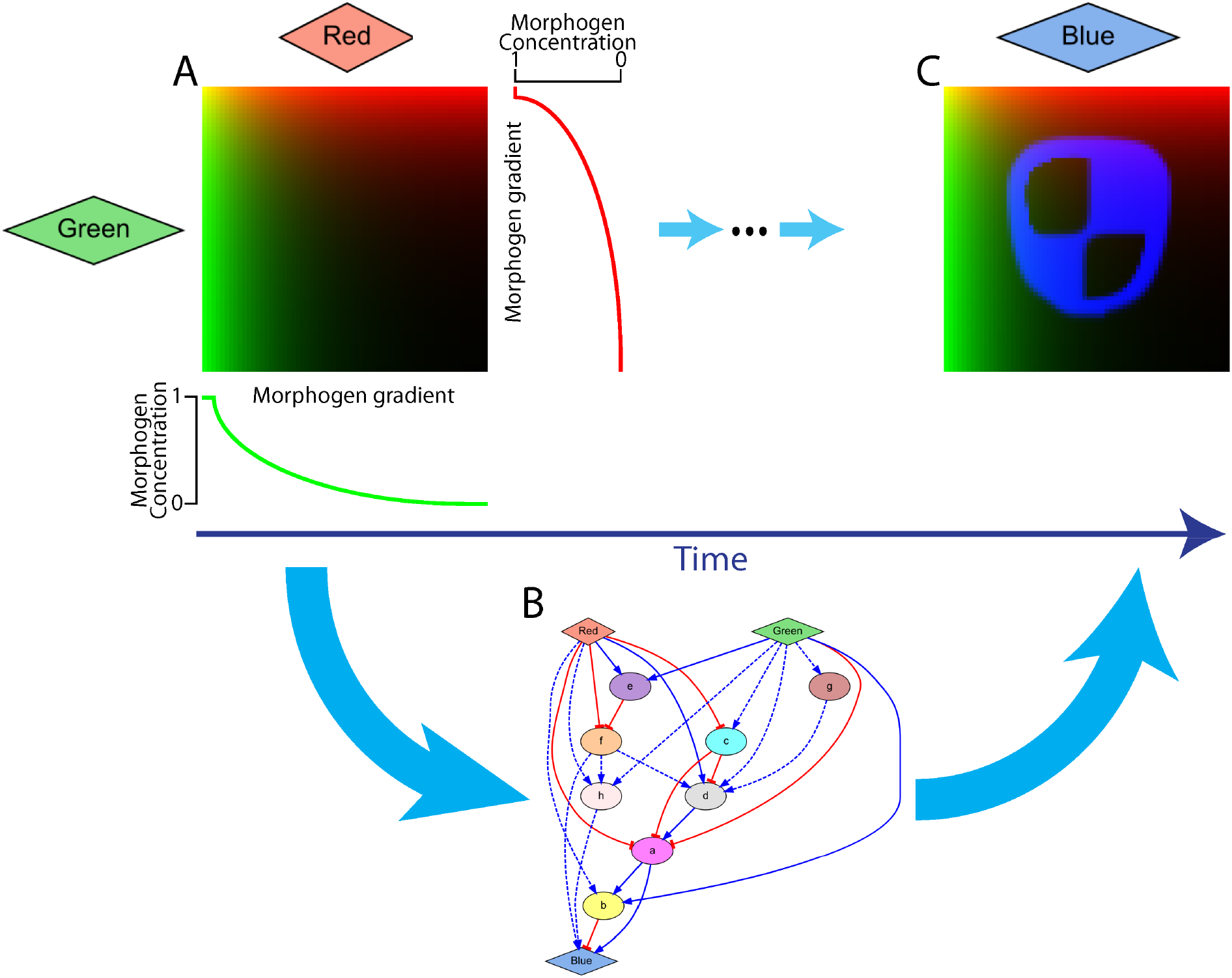
Designing gene regulatory mechanisms for forming a target spatial gene expression pattern in response to two-dimensional orthogonal morphogen gradients. **A.** Two static input morphogens (labeled red and green) form orthogonal gradient signals across a two-dimensional tissue or cell culture. The green morphogen forms a gradient from left to right (left being high), and the red morphogen forms a gradient from top to bottom (top being high). **B.** Each cell in the domain implements the same gene regulatory mechanism (GRM) including the input morphogens, an output reporter gene for the spatial pattern (blue), and intermediate genes (a-h). **C.** The regulatory mechanisms defined in the GRM process the input morphogen gradients so that the expression pattern of the output gene forms the target spatial pattern.

Starting with the input morphogens forming the orthogonal gradients and all the other products at zero concentration, the goal of the designed GRM is to dynamically process the input morphogen signals to express the reporter gene in a stable spatial pattern similar to a target pattern. In addition to the input signals and the reporter gene, the GRM can include intermediate genes to form complex regulatory networks. The input morphogens and intermediate genes can regulate the reporter gene and other intermediate genes, but the reporter gene cannot regulate any other gene. Except the input gradient morphogens, all products are confined intracellularly. GRMs are modeled as a system of partial differential equations (PDEs) where each gene is represented by an equation defining its rate of change in product concentration due to their regulatory interactions and decay.

### 2.2. A versatile modeling framework for non-linear regulatory mechanisms

Biological regulatory interactions between signals and genes are continuous and non-linear and can act as enhancers (positive regulation) or inhibitors (negative regulation). Furthermore, multiple regulatory interactions can affect the same gene in a necessary or sufficient fashion. To allow the design and simulation of such a large variety of gene regulatory mechanisms, we have designed a versatile mathematical approach based on Hill equations combining different terms as building blocks to model the integration of any number of regulatory interactions in GRMs. Single positive and negative regulations follow a sigmoidal response modeled by a simple Hill equation. Multiple positive regulations can be grouped as necessary (similar to an AND gate) or sufficient (similar to an OR gate) by combining multiple terms in the Hill equation numerator with either multiplication or multiplication and summation operators, respectively. For simplicity, negative regulations are always combined with a multiplication operator (AND gate) in this work. Figure 2 illustrates the approach with examples including one and two regulatory interactions. The rate of expression of a product (‘b’) depends on the concentration levels of its regulatory products (‘g’ and ‘r’) as well as the sign (positive or negative) and grouping (necessary or sufficient) of its regulatory interactions. In this way, multiple types of gene regulations can be combined to produce a large variety of continuous regulatory mechanisms, such as the *AND*, *OR*, *NOR*, and *NIMPLY* logic illustrated. Similarly, any number of regulatory interactions can be combined to produce a versatile set of possible genetic mechanisms.

**Figure 2.**
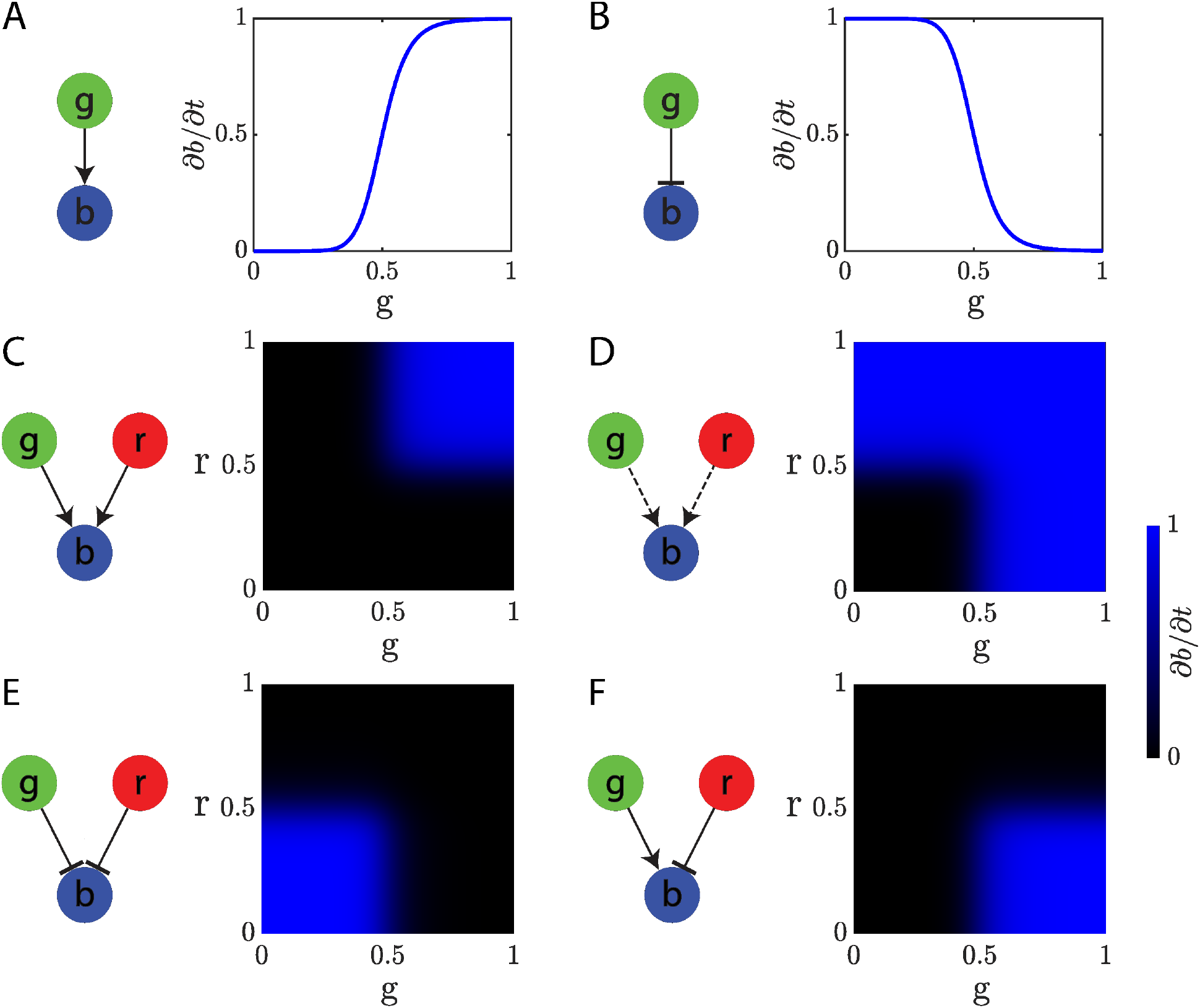
A versatile modeling framework for gene regulatory mechanisms. Using Hill equations combining different terms, continuous non-linear regulatory interactions can be modeled as positive (point arrow) or negative (blunt arrow) and can be grouped as necessary (solid line) or sufficient (dashed line) manner. Illustrative mechanisms are shown with one or two interactions between two input genes (‘g’ and ‘r’) regulating the rate of expression of a reporter gene (‘b’). **A.** Single positive regulation. **B.** Single negative regulation. **C.** Double necessary positive regulation (*AND* logic). **D.** Double sufficient positive regulation (*OR* logic). **E.** Double negative regulation (*NOR* logic). **F.** Double positive and negative regulation (*NIMPLY* logic). Three or more regulatory genes can be combined in a similar fashion to produce complex regulatory interactions.

### 2.3. Automatic design of GRMs

Designing a GRM that can produce a given target function is a current challenge, especially when including spatial features. To streamline this task, we developed a machine learning methodology to automatically design GRMs able to recapitulate the formation of a given spatial pattern. The method makes use of the two-dimensional orthogonal input morphogen gradients together with the versatile regulatory modeling framework based on Hill equations to design GRMs that can form a stable spatial expression pattern. The approach is based on parallel evolutionary computation, where a population of candidate GRMs evolve by iteratively crossing, mutating, simulating, and scoring them until a GRM that can recapitulate the target pattern is found (Fig. 3). The method takes as input a target spatial pattern (e.g., a square shape) and returns a complete GRM—including the number of intermediate genes, regulatory interactions, and parameters—that when simulated produces a reporter gene with a spatial expression pattern similar to the given target pattern.

**Figure 3.**
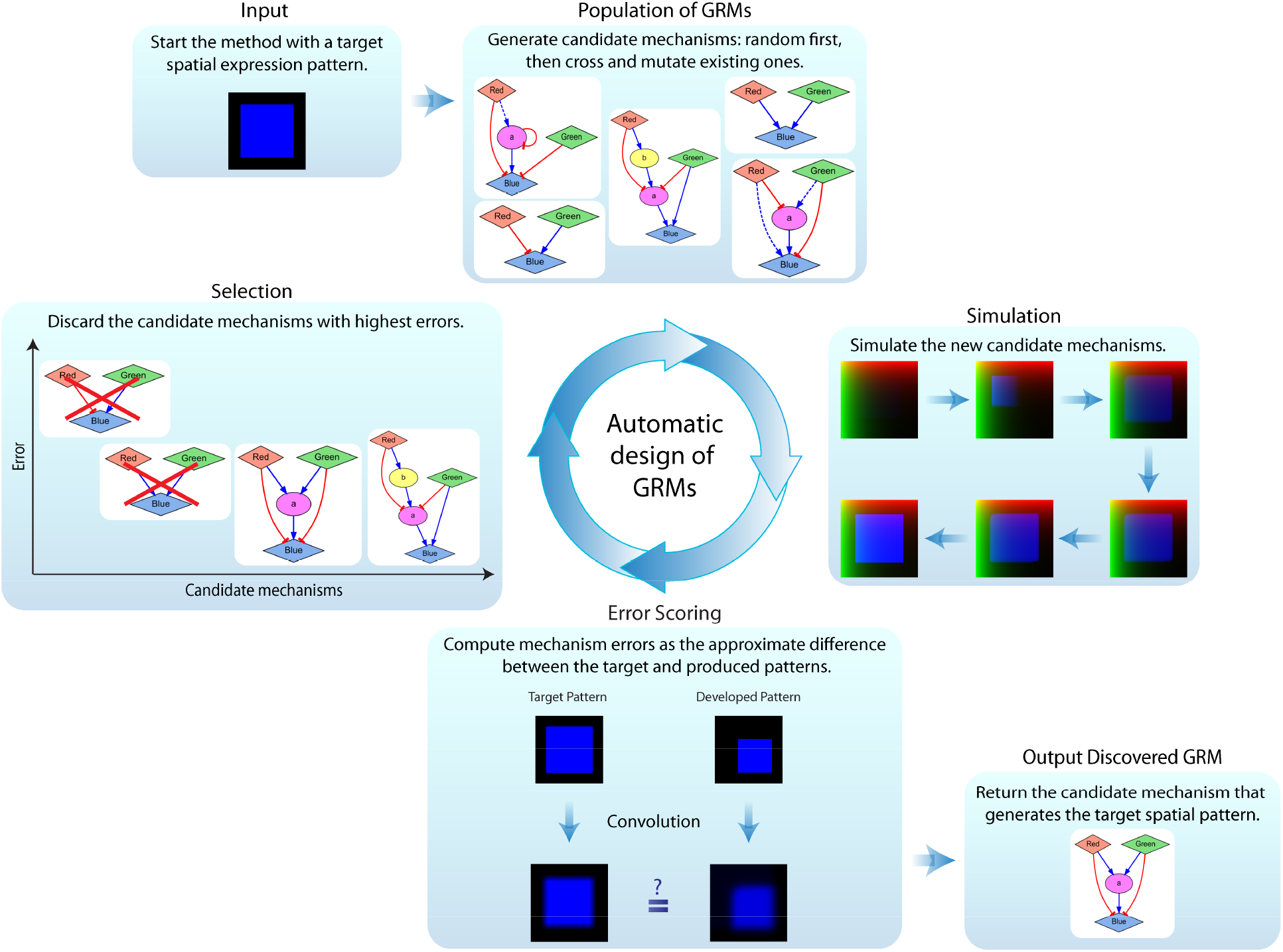
Automatic methodology based on evolutionary computation for the design of gene regulatory mechanisms producing a given spatial expression pattern. The method takes as input a target spatial pattern (a square shape in this example), iteratively generates a population of candidate mechanisms by crossing and mutating them, simulates and computes their errors as their ability to produce the target pattern, and discards those with the highest errors. This iterative process repeats until a regulatory mechanism with zero error is discovered.

The pseudocode for the evolutionary algorithm to design regulatory mechanisms is described in Box 1. The algorithm starts with a random population of GRMs, each including the gradient input signals and the reporter gene, a random number of intermediate genes, random interactions among all products (except the gradient signals, which have no regulators, and the reporter gene, which cannot regulate other products), and random parameters. Based on the presented framework for non-linear regulatory mechanisms, GRMs are translated into a system of partial differential equations, which can then be numerically simulated to score their capacity to produce the target expression pattern (error of the model). The GRMs that produce the most similar and stable patterns as compared to the input target pattern are kept in the population, while GRMs with higher errors are discarded. The population then produces new offspring GRMs by stochastically crossing those in the current population and adding random mutations. The new offspring GRMs are then simulated, scored, and added to the population to select the next generation. This iterative process continues until a GRM with zero error is found, representing a GRM that can produce the input target pattern.

##### Box 1. Evolutionary algorithm pseudocode for the design of GRMs

1 Initialize population with random GRMs
2 Simulate GRMs
3 Compute fitness of GRMs by comparing the developed and the input target patterns
4 While (best GRM fitness > 0)
4.1 Reproduce population of GRMs by crossover and mutation
4.2 Simulate children GRMs
4.3 Compute fitness of children GRMs by comparing the developed and the input target patterns
4.4 Select best GRMs for next generation population
5 Return best GRM found

The fitness of a GRM scores its ability to stably form the target spatial pattern. It is computed at the last time step of the simulation as the sum of the average difference between each domain location (pixel) in the target and the developed pattern plus the maximum concentration change (penalizing patterns not in equilibrium). Thresholds are defined for both measurements to avoid overfitting (parameters *α* and *β*, respectively). In addition, the spatial pattern fidelity required for the designed GRMs can be adjusted in the method, since sharp and complex pattern boundaries may require excessively complex GRMs. Before calculating the fitness, both the target and developed patterns are processed with a box blur kernel convolution to eliminate sharp spatial features. Similar to the concentration and equilibrium thresholds, the strength of the convolution function can be adjusted with a parameter defining the kernel size (parameter *k*; higher values representing more approximate patterns). In this way, the fidelity of the GRM to produce the target pattern can be adjusted with error parameters at the concentration, equilibrium, and spatial levels.

### 2.4. Inferred GRMs for geometric patterns

We tested the proposed methodology for the design of GRMs that can produce different geometric target patterns. Figure 4 shows an illustrative example of the evolutionary dynamics of the algorithm across three independent runs for the design of a GRM to develop a spatial pattern with a triangle shape. The initial models are random and produce patterns far from the target, hence with high error scores. After many generations of crossover and mutations, including adding *de novo* intermediate nodes, the models increase their complexity in terms of number of genes and links and their ability to produce the target spatial pattern. After ∼5,000 generations taking ∼20 hours of computation time, the evolutionary process finds GRMs that can produce the target triangle pattern with zero error (for the given error thresholds).

**Figure 4.**
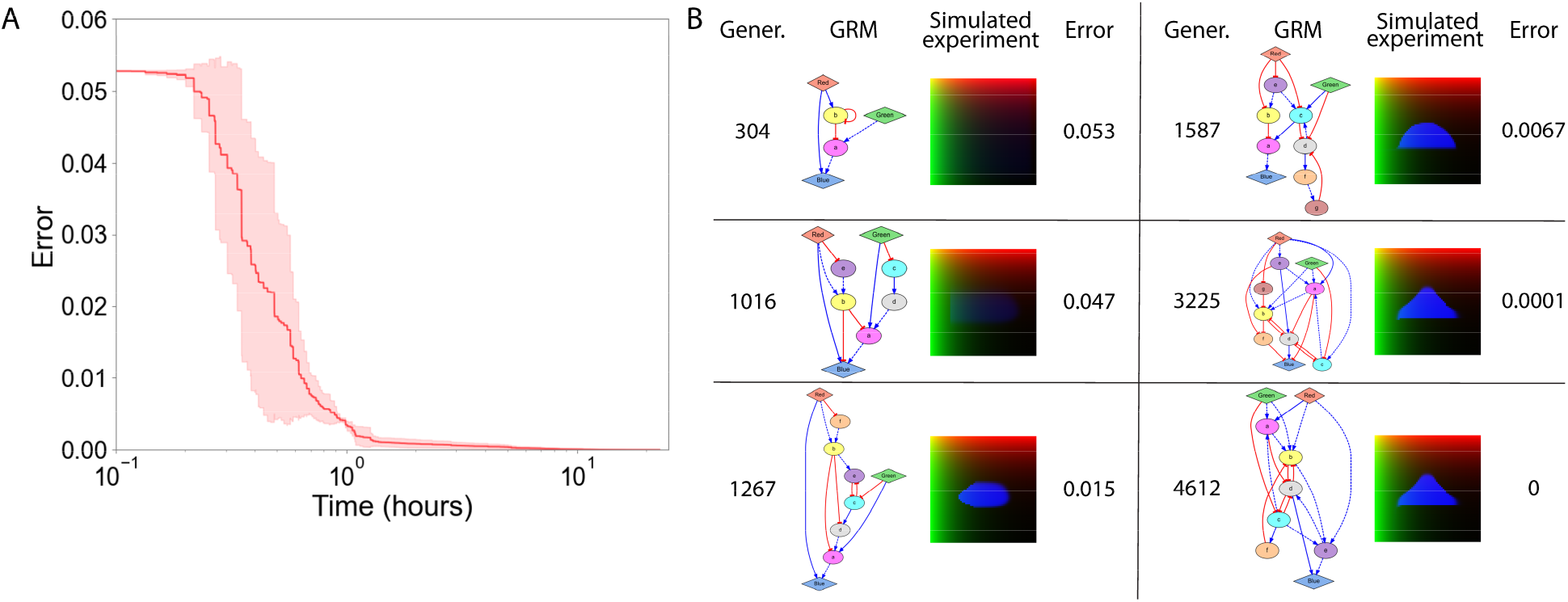
Evolutionary dynamics for the design of a GRM producing a triangle pattern. **A.** Average error of the best GRM across three independent runs of the algorithm, all reaching zero error after ∼20 h of execution. The shaded area represents the standard deviation. Error parameters: *k* = 5, *⍺* = 0.1, *β* = 0.001. **B.** Representative candidate GRMs at different generations during the evolutionary search, showing the gradual increase in complexity and improvement in their capacity to produce the target triangle pattern.

The regulatory links and intermediate genes automatically added by the evolutionary algorithm perform the necessary spatial computations to produce the target spatial patterns. Figure 5 shows examples of the resultant GRMs discovered by the automated methodology for four different target geometric shapes, all found in less than 35 hours of computational time (Supplementary Fig. 1 and Supplementary Movies). Target patterns with edges parallel to the orthogonal gradients (Fig. 5A; square pattern) require less complex GRMs than patterns with oblique lines (Fig. 5C-D; triangle and diamond). To produce curved edges (Fig. 5B, circle), the algorithm takes advantage of regulatory interactions with gradual slopes, which produce softer responses at the edges.

**Figure 5.**
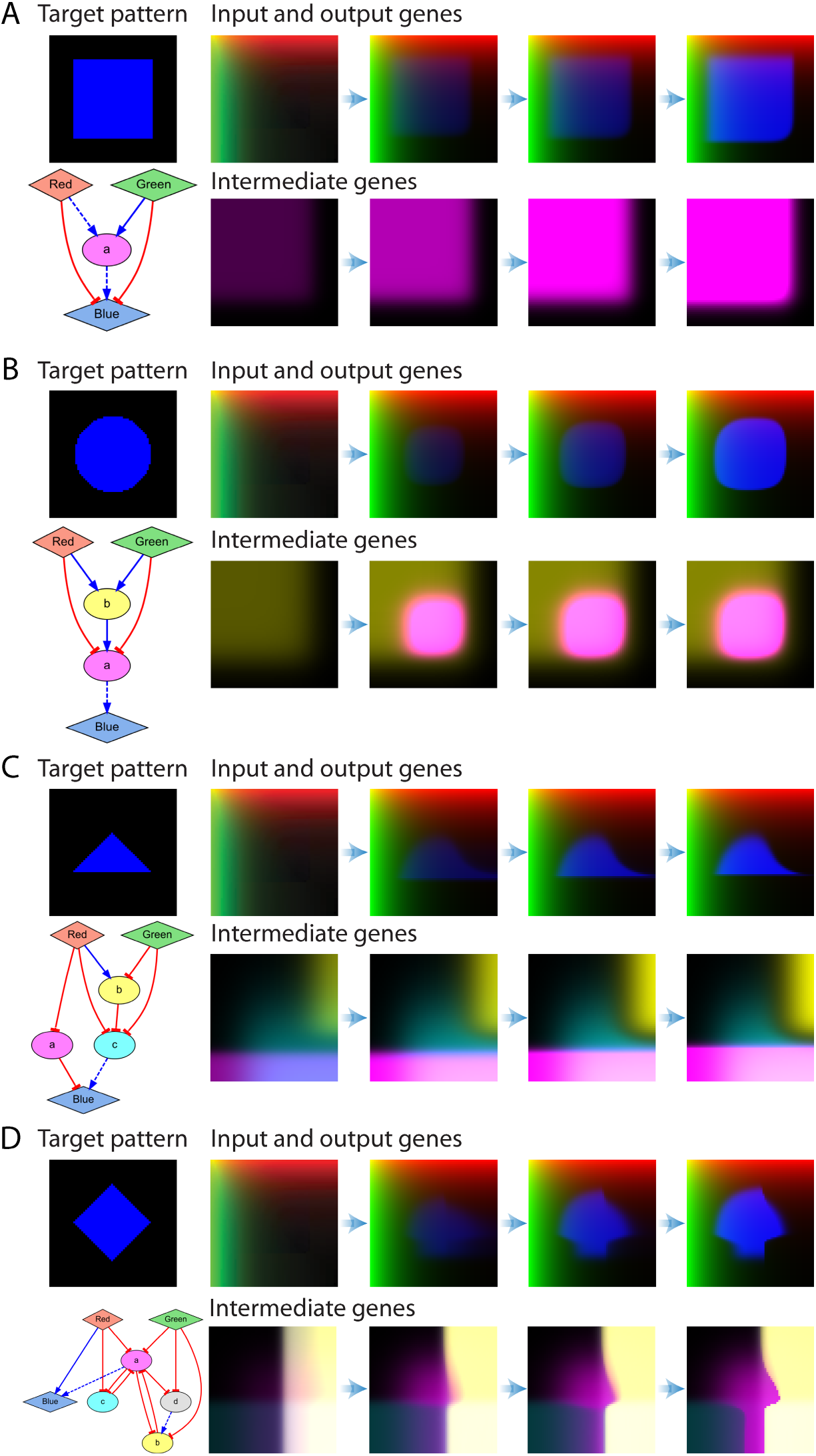
Automatically designed GRMs for different geometric target shapes. The simulation of a GRM starts with all the genes at zero concentration except the input signals (green and red), which form static orthogonal gradients. All the resulting GRMs correctly produce a steady-state target pattern (blue) with zero error. The dynamics of the output and intermediate genes reveal the spatial computations performed by each mechanism to produce their target patterns. Expression colors in the simulation correspond to the node colors in the GRM network diagrams. Error parameters: *k* = 7, *⍺* = 0.25, *β* = 0.001.

The simulation dynamics show how the regulatory interactions designed in the GRMs translates into spatial computations to produce the target patterns. For the square pattern (Fig. 5A), the negative regulations between the input gradient signals (red and green) and the output reporter gene (blue) prevent the latter from being expressed at high input concentrations (top and left sides of the domain). In addition, the input signals positively regulate with a low threshold an intermediate gene (‘a’), resulting in no expression in the bottom and right areas. Then, a positive regulation between the intermediate and output gene defines the bottom and right edges of the square pattern. The GRM for the circle pattern (Fig. 5B) extends this design with an additional gene (‘b’) at the end of the pathway that together with more gradual regulatory interactions produce the curved edges needed for the circle pattern. The discovered GRMs for the triangle and diamond patterns (Fig. 5C-D) include three and five intermediate genes, respectively, that define intermediate expression patterns at different domain locations to produce the target geometric shapes.

### 2.5. Adjusting the complexity and pattern precision of the designed GRMs

The tolerance of the proposed method can be adjusted for different design needs, from complex regulatory mechanisms that produce exact patterns to simple mechanisms that produce approximate patterns. For this, different values can be set for the kernel size (*k*) and concentration threshold (*α*) parameters used in the error fitness function. Figure 6 shows the results of lowering these parameters to design more complex GRMs for the same geometric target shapes as for the previous simpler networks, all with zero error. While the simpler GRMs produced approximate patterns with diffuse edges (Fig. 5), especially those that are not parallel to the gradient signals, the complex GRMs produced precise patterns with sharp edges, even at different angles to the gradient signals or forming curves (see also Supplementary Movies). Conversely, the time needed by the algorithm to design complex networks that can produce precise patterns was significantly longer (about 5x) than for designing simpler networks to produce approximate patterns (Fig. 6E).

**Figure 6.**
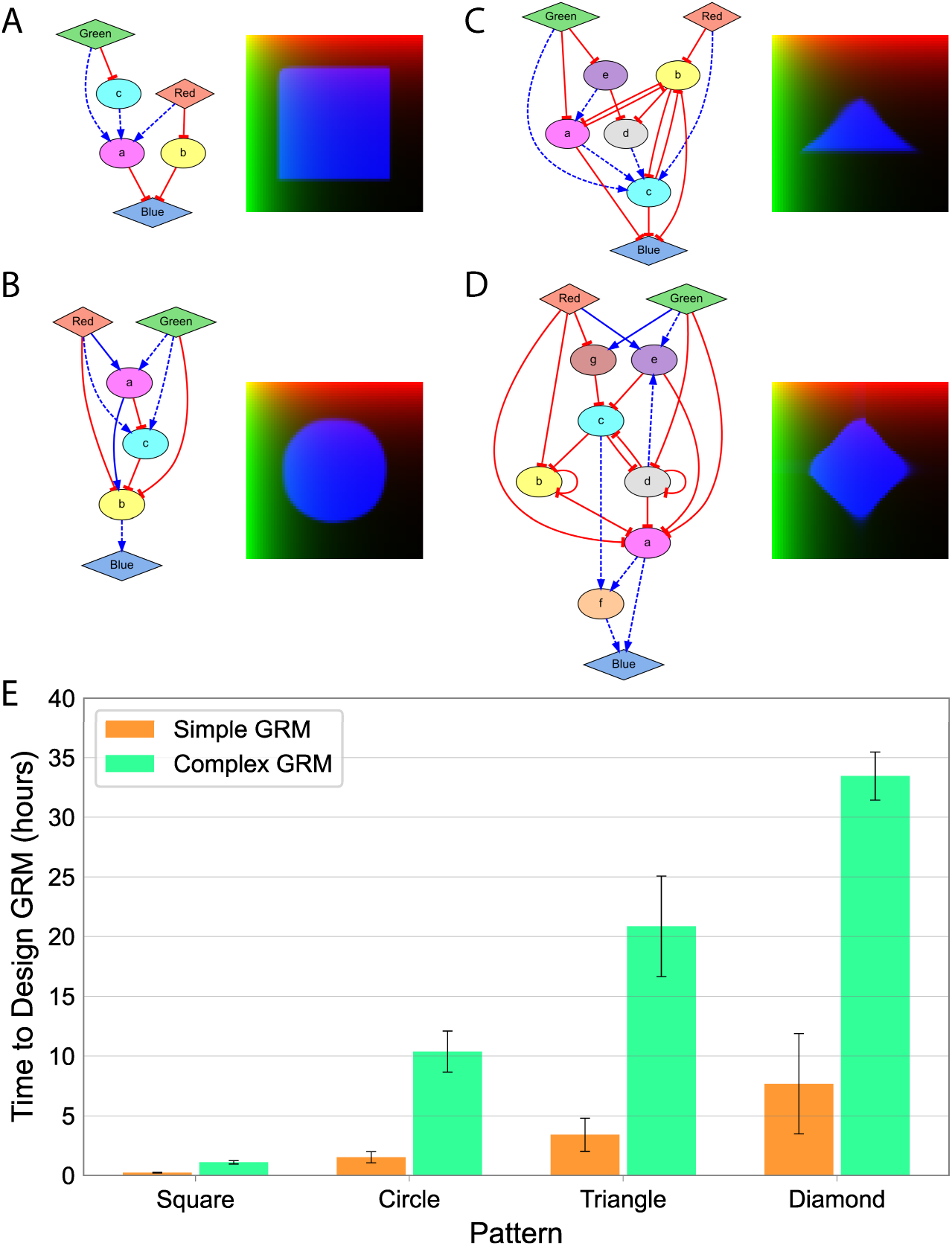
The method includes tolerance parameters to control the complexity of the designed GRMs. The kernel size (*k*) and error concentration threshold (*⍺*) parameters determine the complexity of the designed GRMs and their precision in producing the target patterns. **A-D.** Lowering the tolerance parameters results in more complex GRMs but with more precise patterns as compared to Fig. 5. Error parameters: *k* = 5, *⍺* = 0.1, *β* = 0.001. **E.** Time needed by the algorithm to design simple (Fig. 5) or complex (A-D) GRMs with zero error. Each condition was run three times. Error bars denote standard deviation.

The complexity of the GRMs designed consistently depends on the target pattern and the tolerance parameters used. Figure 7 shows the average complexity of GRMs for the square, circle, triangle, and diamond target patterns obtained with different values of kernel size and error concentration thresholds. The results illustrate how the complexity of the discovered GRMs for each pattern decreases as the kernel size and the error concentration threshold parameters increase. The effect of these parameters is more acute in the case of complex patterns (Fig. 7D) as compared to simpler ones (Fig. 7A). Moreover, increasing the error concentration threshold results in more diffuse edges in the developed patterns, while increasing the kernel size results in less precise shapes (Supplementary Figs. 2-5).

**Figure 7.**
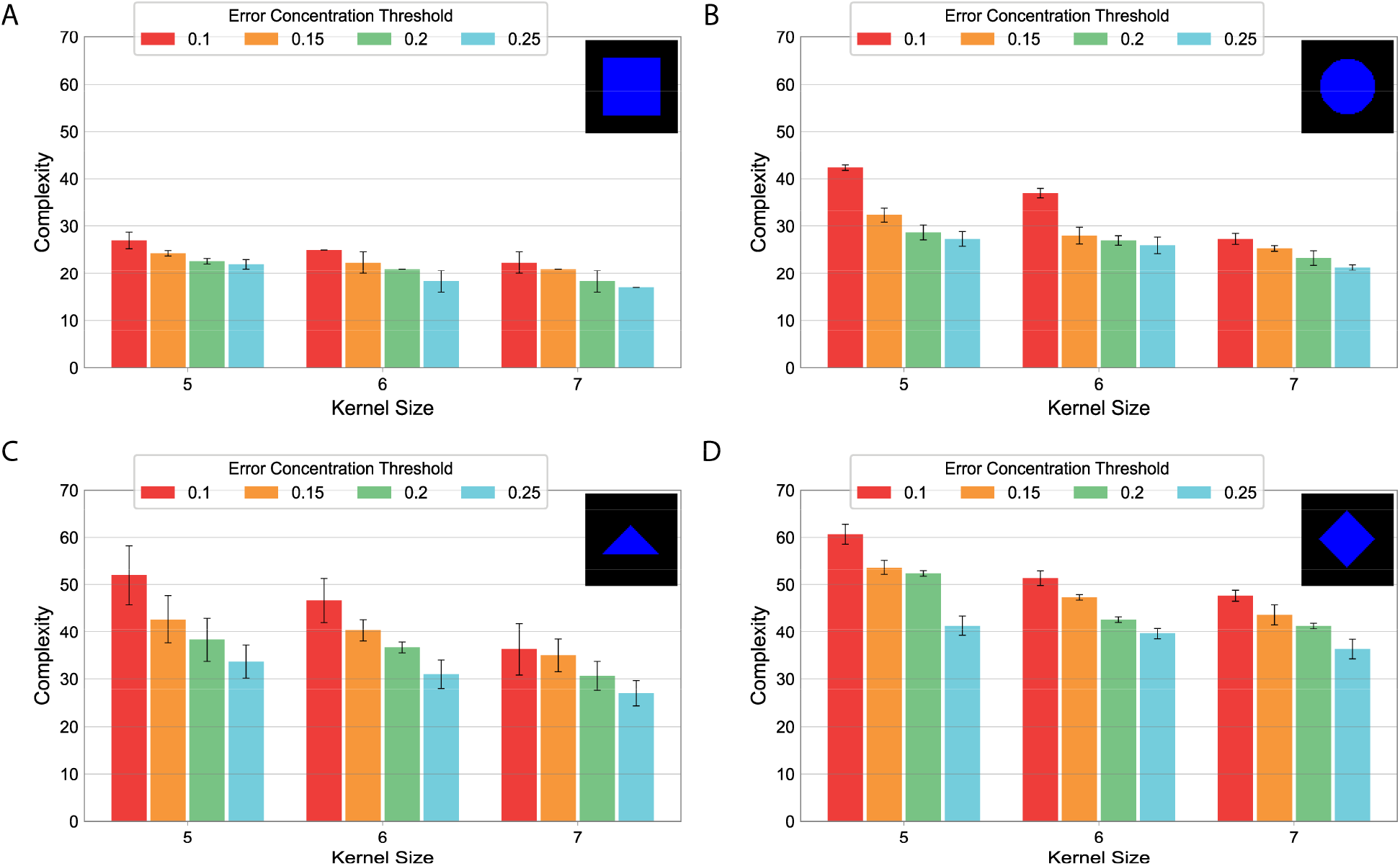
Complexity of the designed GRMs discovered by the automated search method for the geometric shapes with different tolerance parameters. The complexity of a GRM was measured as the number of links plus three times the number of genes. Three runs were performed for each target pattern, including the square (A), circle (B), triangle (C), and diamond (D), and parameter values for kernel size (*k*) and error concentration threshold (*α*). Error bars denote standard deviation.

### 2.6. Designing GRMs for arbitrary shapes

To test the ability of the method to design GRMs for arbitrary shapes, we applied it to discover models that can produce gradients, periodic shapes, symbols, and characters. Figure 8 shows the developed expression patterns produced by the designed GRMs, all reaching zero error (see Supplementary Figs. 6-7 and Supplementary Movies for the target patterns and the expression of intermediate genes). The complexity of the discovered networks varies from 29 to 91 (as the number of edges plus three times the number of genes). The results show how the method can design GRMs for complex shapes with fine details such as gradients, curved lines, and pointed ends—all produced by the interpretation of two orthogonal morphogen gradients (red and green).

**Figure 8.**
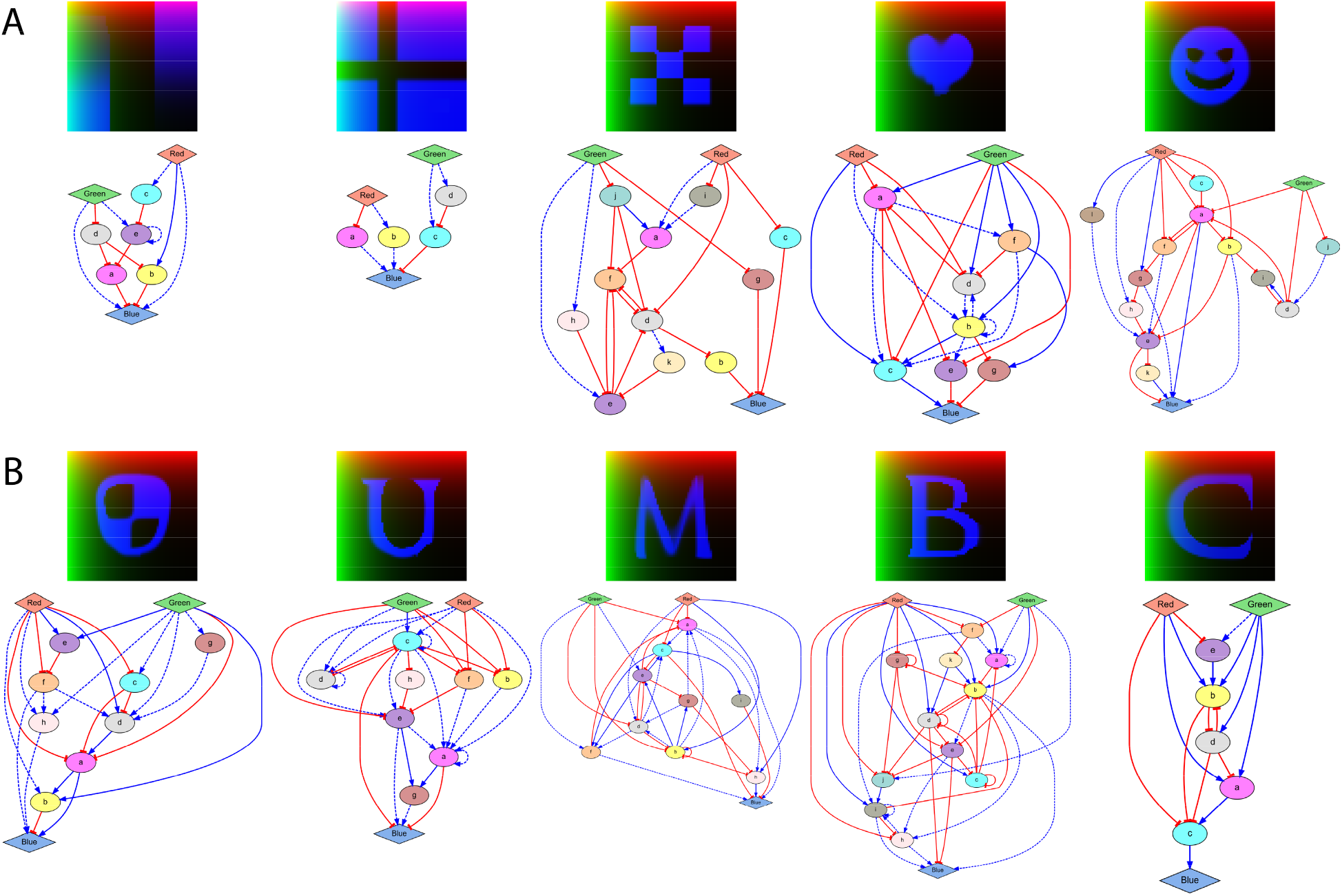
Patterns produced by GRMs discovered by the automated methodology applied to arbitrary shapes. The patterns tested for arbitrary shapes include: (**A**) gradients, flag, checkerboard, and symbols; (**B**) logo and characters. Parameters: *k* = 5, *⍺* = 0.1, *β* = 0.001.

## 3. Discussion and Conclusions

To streamline the design and understanding of gene regulatory mechanisms (GRMs) capable of producing spatial patterns in response to morphogen gradients, here we proposed a novel methodology integrating a framework to model arbitrary genetic circuits with evolutionary computation and high-performance computing. The method can design synthetic GRMs— including all the necessary genes, regulatory links, and parameters—that when simulated develop a given target spatial pattern in response to orthogonal morphogen gradient signals. A GRM is formalized as a PDE system, which is generated from a parallel implementation of a population-based heuristic optimization algorithm. We demonstrated the capacity of the method to design GRMs able to interpret the positional information provided by the signal gradients and produce a variety of target patterns, including geometric shapes, symbols, and characters. Crucially, the precision of the designed GRMs to produce a given pattern can be adjusted with a convolution-based fitness function that evaluates their ability to recapitulate the target pattern. The results showed how these parameters can modulate the trade-offs between the precision of the produced pattern, the complexity of the designed mechanism, and the speed of the machine learning method. Hence, while the presented methodology can design very complex GRMs, it is also versatile enough to automatically limit the complexity of such networks for a variety of applications and studies.

The proposed methodology is highly versatile but could be extended with further functionality in future work. The products of intermediate genes are restricted to intercellular pathways, but the methodology could include intermediate genes that produce diffusive signals, such as ligands acting intracellularly. This would allow more complex spatial regulation involving dynamic morphogens, as found in developmental processes (Dickmann et al., 2022) and novel synthetic biology applications (Oliver Huidobro et al., 2022). We have limited this study to GRMs with only one reporter gene, but the method could be used to design GRMs with multiple outputs to produce target patterns involving multiple gene products. Similarly, the framework is currently limited to two-dimensional domains and target patterns, but it could be easily expanded to three-dimensional pattern systems (Sohka et al., 2009). The machine learning method can explore and use a large range of *in silico* regulatory interactions and degradation rates. However, the method could be restricted to use a predetermined set of standard biological parts to facilitate the engineering of the automatically-designed GRMs into synthetic biological systems (Bird et al., 2022; Buecherl and Myers, 2022; Martínez-García et al., 2023). We have assessed the performance of the method with the design of GRMs for synthetic patterns and shapes, but it could be applied to the understanding of patterning in developmental biology. Using as input a target pattern found in a natural biological system, the method could be used to generate hypotheses in the form of GRMs recapitulating the observed spatial phenotypes (Lobo and Levin, 2015) that subsequently could be validated experimentally (Lobo et al., 2016). Finally, the algorithm is limited to return a single GRM for each run. However, multiple GRMs could lead to the formation of the same pattern. Indeed, a major challenge for complex phenotypes is to discover a comprehensive set of GRNs that can develop a given gene expression pattern, i.e., an atlas of regulatory designs (Cotterell and Sharpe, 2010; Scholes et al., 2019). Future work will extend the presented methodology with evolutionary multi-objective and diversity-preserving algorithms (Liu et al., 2017) for the discovery of a comprehensive set of GRMs that can produce a given pattern.

In conclusion, the capacity of the presented method to automatically design GRMs for spatial patterns could be essential to transition from our current ability to understand and implement synthetic small modules to be able to identify and assemble larger scale systems (Purnick and Weiss, 2009). Recent advancements have made it possible to streamline the engineering of synthetic GRMs for a given arbitrary circuit (Jones et al., 2022). The presented methodology expands these approaches by automating also the design of such circuits for producing complex patterns, which could potentially be applied to current bioengineering problems—from the synthesis of complex bioproducts in industrial, pharmaceutical, and biomaterial applications (Hwang et al., 2020) to the construction of multicellular synthetic systems (Davies and Levin, 2023). In addition, these advancements could streamline the systematic study of natural circuits towards the discovery of complex mechanisms controlling tissue spatial behaviors (Ko and Lobo, 2019) and whole-body patterns (Herath and Lobo, 2020) from morphogen gradients. Indeed, it is a current challenge to produce mechanistic hypotheses in terms of GRMs that can recapitulate observed spatial phenotypes. The methodology presented here could be employed to automatically infer such hypotheses directly from datasets of curated experimental gene expression patterns (Lobo, 2022; Roy et al., 2020). Overall, the presented method to aid in the design of GRMs able to produce arbitrary gene expression patterns has the potential to both enable the understanding of complex developmental processes as well as the design of complex dynamic synthetic systems.

## 4. Methods

### 4.1. Simulation of GRMs

We developed a simulator of GRMs for spatial pattern formation based on a system of nonlinear partial differential equations (PDEs). Gene regulations are based on Hill equations and a GRM consists of two input morphogens, intermediate genes, and a reporter gene. The two input morphogens (red *R* and green *G*) have a constant gradient distribution given by *M*_R_ = *d*^j^, *M*_G_ = *d*^i^, where *d* = 0.93 and (*i*, *j*) is the cell position in the domain. Intermediate genes can be activated or repressed by other genes except for the output gene, which can be regulated but cannot regulate other genes—since it represents a reporter signal that defines the output pattern produced by the GRM. Gene products are confined intracellularly and decay over time.

Each gene in a GRM includes four parameters: production, decay, diffusion, and basal expression. Each regulatory interaction is modeled as positive (activating) or negative (inhibiting) using Hill equations and include two parameters: Hill coefficient and binding constant. Genes can be regulated by several other genes simultaneously, and these regulations can be grouped as necessary or sufficient. Necessary positive regulations are combined with a multiplication operator (AND logic), while sufficient positive regulations are combined with both a multiplication and summation operation (OR logic). Negative regulations are combined with a multiplication operator (AND logic). This methodology guarantees that the strength of any regulation lies within the range [0,1]. The illustrative examples of single and double regulations shown in figure 2 are modeled by the following equations:

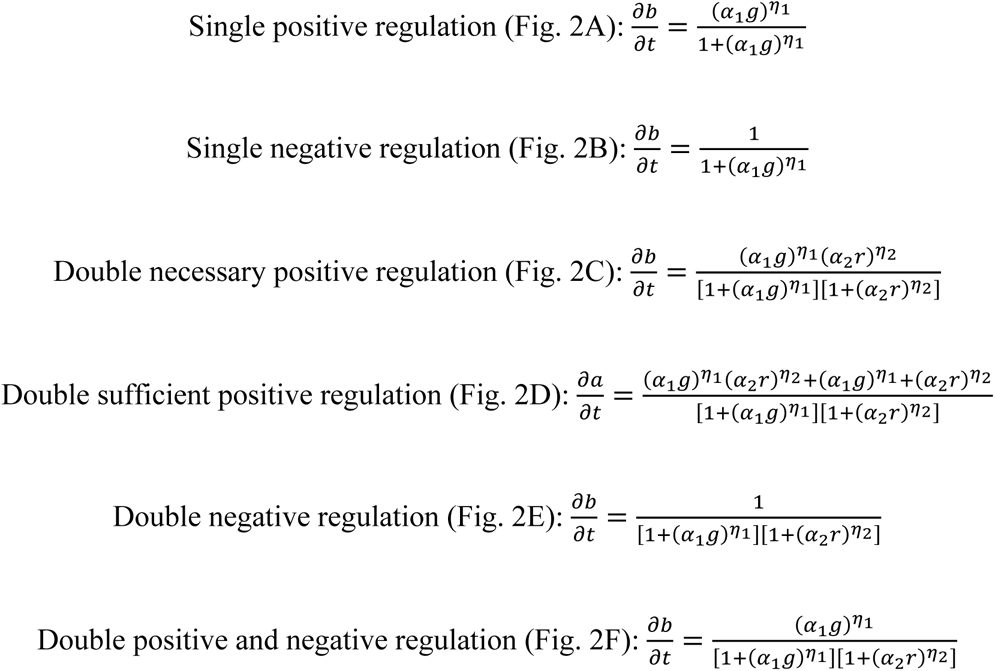

where *η*_i_ is the Hill coefficient and *⍺*_i_ is the binding constant in the regulatory interaction *i* (*η*_i_ = 10 and *⍺*_i_ = 2 for all regulations shown in Fig. 2).

In this way, the rate of change of a gene can integrate any combination of interactions. For example, the following PDE describes a gene *a* regulated by two activators *b* and *c* as sufficient, two activators *d* and *e* as necessary, and an inhibitor *f*:

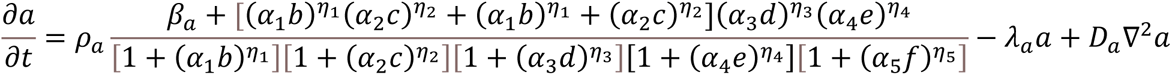

where *ρ*_a_is the gene production constant, *β*_a_is the gene basal expression level, *η*_i_are the regulation Hill coefficients, *⍺*_i_ are the regulation binding constants, *λ*_a_ is the gene decay constant, and *D*_a_ is the gene diffusion constant. All parameter units are arbitrary. The basal expression level is zero when a gene receives at least one positive regulation; otherwise, it is one to model a constitutive promoter that can be inhibited by other genes.

### 4.2. GRM fitness error

The error of a developed expression pattern is calculated at the last time step of the simulation (100 steps with *dt* = 1 in this work) by comparing it to the input target expression pattern with a kernel-based method. In this approach, both the developed and target patterns are first blurred by a box blur kernel convolution. The error of a candidate GRM is then calculated as the sum of the average log difference between each domain location (pixel) in the simulated pattern from the candidate GRM and the target pattern, plus the maximum concentration change at the last time step, which penalizes patterns not in equilibrium. In this way, the fitness function represents an approximate comparison between the developed and target images, ignoring small details. Hence, the error between a developed expression pattern *D* at the last time step in the simulation and the input target expression pattern *T* is calculated as:

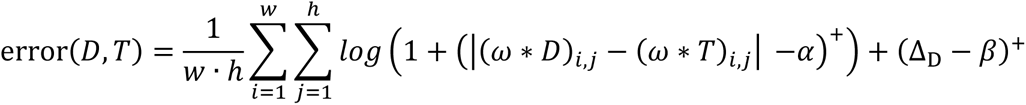

where *w* and ℎ are the simulation domain width and height, respectively, *k* is the kernel size, *ω* is the box blur kernel defined as 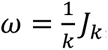, where *J*_k_ is the unit matrix, *α* is the error concentration threshold, Δ_D_ is the maximum concentration change in *D* at the last time step in the simulation, and *β* represents the equilibrium penalty threshold. (*x*)^+^indicates the positive part function of x, which outputs 0 if it is negative and *x* if x is nonnegative. *f* ∗ *g* represents the convolution of *f* and *g*.

### 4.3. Machine learning method

The machine learning method is based on evolutionary computation to automatically design a GRM that can produce a given pattern. The algorithm evolves a population of candidate regulatory mechanisms iteratively by reproduction, fitness calculation, and selection to find an optimal GRM. The population follows an island distribution approach (Whitley et al., 1999) to maximize parallelism and ensure robustness and diversity of the candidate mechanisms. The method produces new GRMs by stochastically mixing existing ones and adding random mutations in each generation.

A crossover creates two new children GRMs by randomly combining two existing regulatory mechanisms in the population. For this, the genes of both parents along with their regulatory links are distributed randomly to the two children GRMs (without gene duplication or changes in kinetic parameters). Next, mutations are applied randomly so that genes and regulations can be added, removed, and their parameters replaced with new ones within their ranges from a random uniform distribution. The input and output genes cannot be removed. Deletion mutations were set to a higher probability than duplication mutations to bias the algorithm towards simpler mechanisms and prevent bloating (Luke and Panait, 2006). A deterministic crowding method (Mahfoud, 1992) was used to select new offspring when their errors (fitness) are equal or better than their closest parents. The algorithm runs until a GRM with zero error is found and 250 additional generations have passed without a decrease in its complexity (number of edges plus three times the number of genes of the simplest GRM with zero error).

The parameter ranges for the mutation operator uniform distributions were set as follows: Hill coefficient (1,10), binding constant (1,100), decay constant (0.1,1), and production constant (0, 0.1), except for the input morphogens, which is set to 0. Diffusion constants were set to zero for all products, in which case they are considered exclusively intracellular. The meta-parameters of the machine learning method were set as follows: crossover rate 75%, mutation rate 1%, link/gene duplication rate 1%, link/gene deletion rate 7.5%, equilibrium penalty threshold 10^-3^. All search runs used 32 subpopulations (islands) with 64 individuals each. Islands are randomly paired, and their regulatory mechanisms are shuffled every 250 generations.

### 4.4. Implementation

The methodology was implemented in C++ with the standard, Eigen (Gaël Guennebaud, Benoît Jacob, and others), Qt (The Qt Company Ltd.), and Qwt (Uwe Rathmann and Josef Wilgen) libraries. We implemented the Euler finite difference method (Press et al., 1986) to numerically solve the system of PDEs in a 64×64 domain (see Supplementary Information for the equations for all the GRMs presented). The method used 48 parallel threads and was run on a computer server with two 24-core Intel Xeon Gold 6240R CPUs at 2.4 GHz and 192 GB DDR4 RAM to evaluate its performance. The source code with the method is freely available in GitHub (https://github.com/lobolab/grm-design).

## Acknowledgments

We thank the members of the Lobo Lab for helpful discussions. This work was supported by the National Institute of General Medical Sciences of the National Institutes of Health under award number R35GM137953. The content is solely the responsibility of the authors and does not necessarily represent the official views of the National Institutes of Health. Computations used the UMBC High Performance Computing Facility (HPCF) supported by the NSF MRI program grants OAC-1726023, CNS-0821258, and CNS-1228778, and the SCREMS program grant DMS-0821311.

