## Supplementary Information for "Automatic design of gene regulatory mechanisms for spatial pattern formation"

This supplement includes additional figures and the system of equations for each of the GRMs presented in the paper.

#### 1. Supplementary Figures

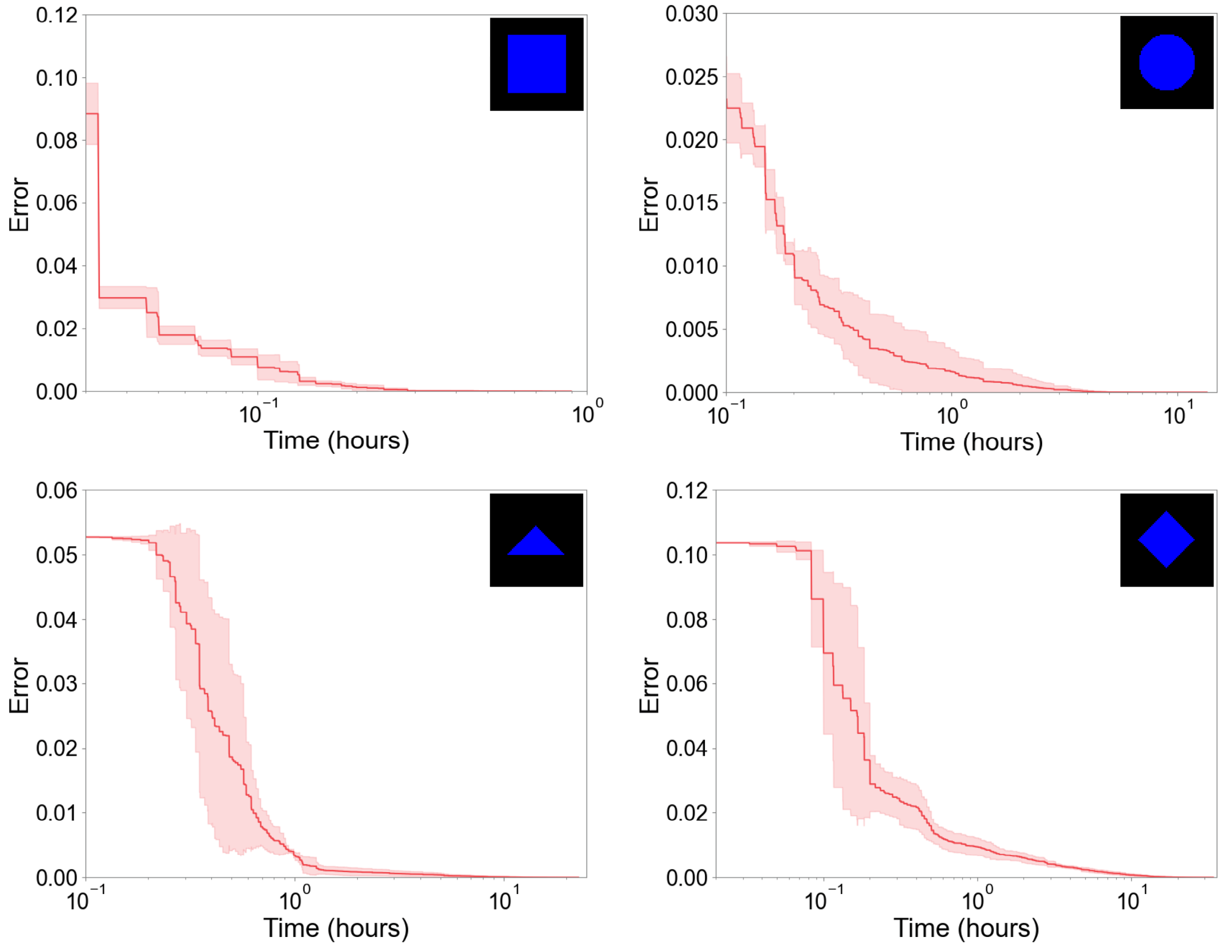


**Supplementary Figure 1. Evolutionary dynamics for the design of GRMs producing geometric target shapes.** Average error of the best GRM across three independent runs of the algorithm, all reaching zero error. Each plot has a different y-axis and x-axis range for better visualization. The shaded area represents the standard deviation. Error parameters: $k = 7$, $\alpha = 0.25$, $\beta = 0.001$.

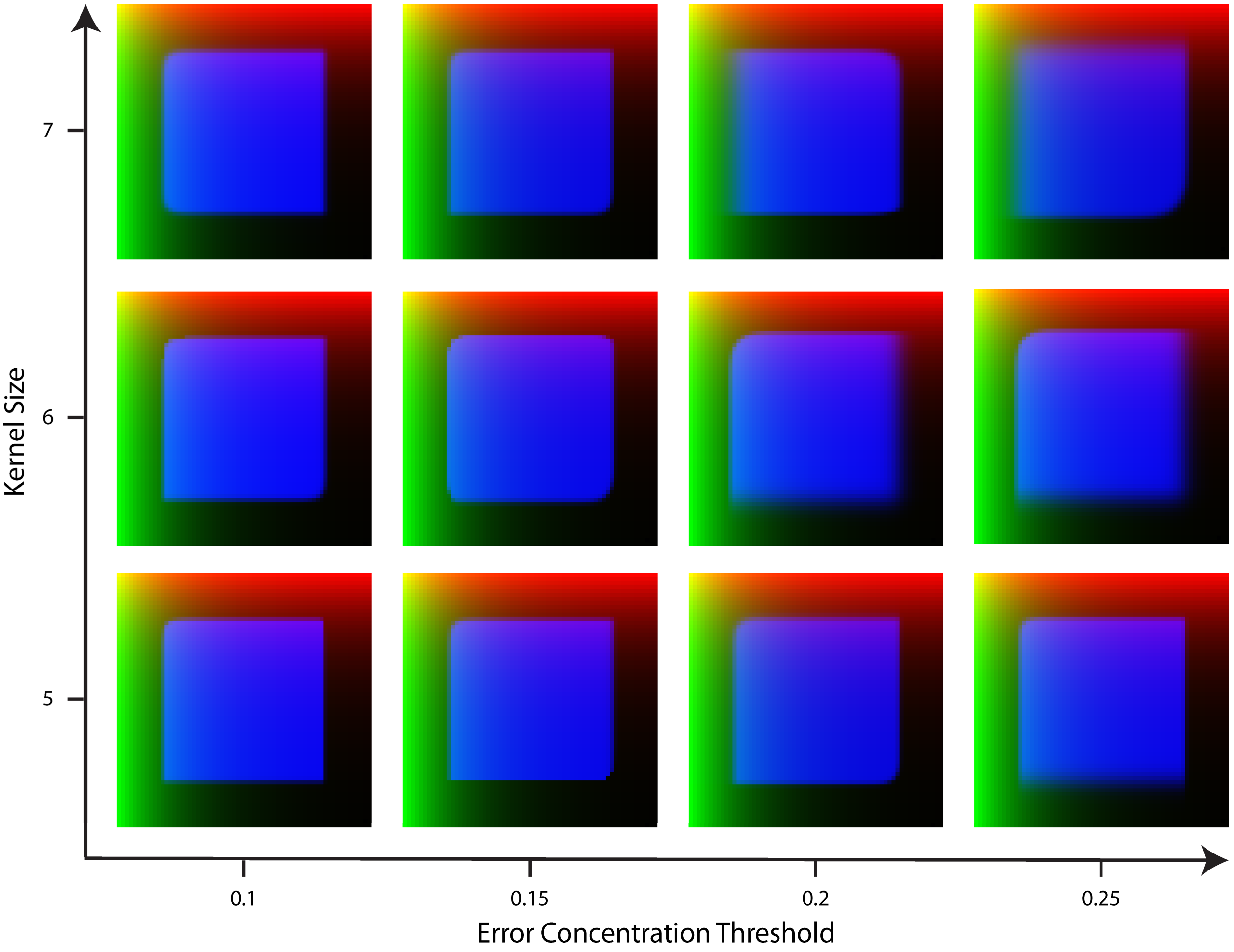


**Supplementary Figure 2. Patterns developed by GRMs discovered with the automated methodology for a square target pattern using different values of kernel size and error concentration thresholds.**

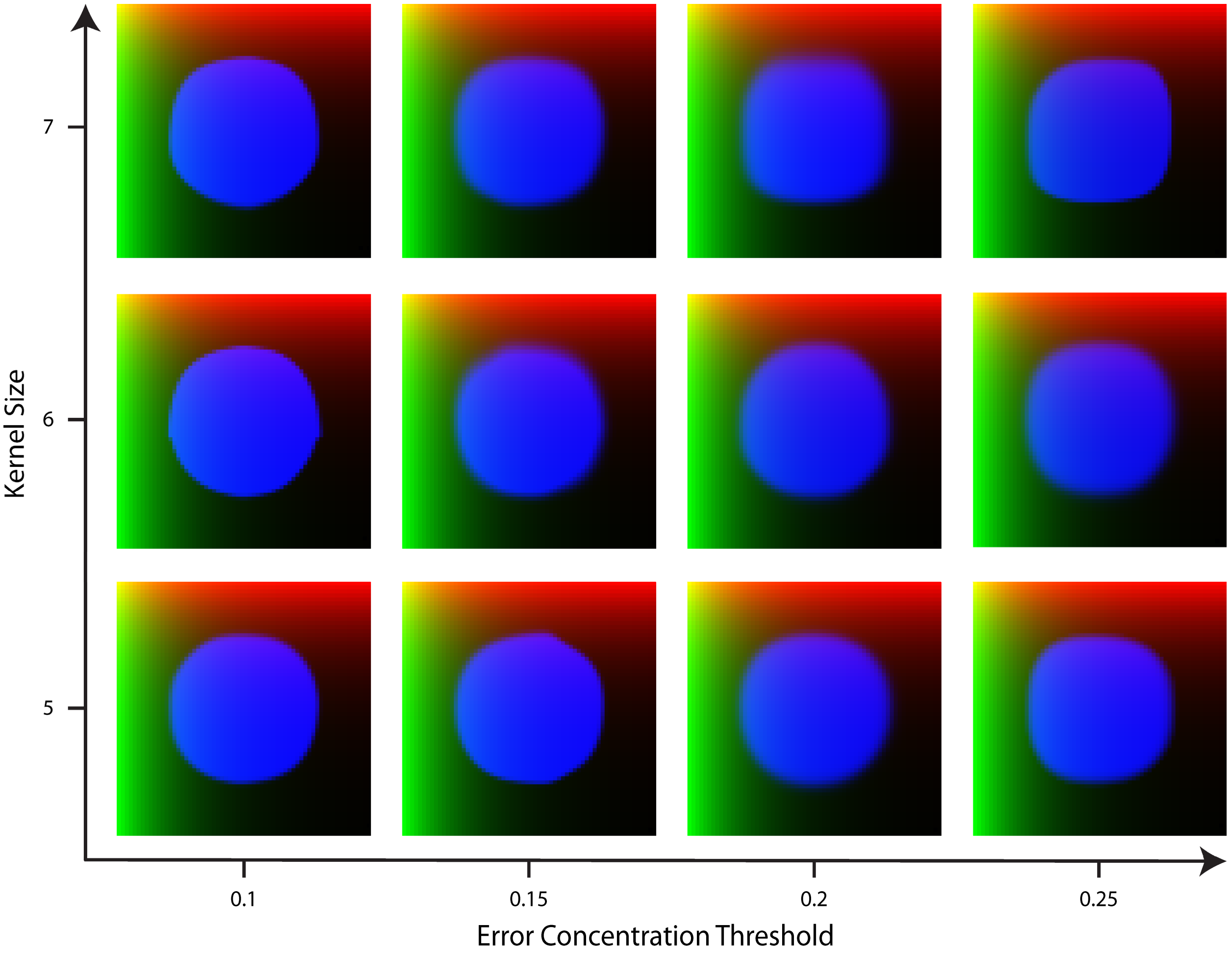


**Supplementary Figure 3. Developed patterns by GRMs discovered with the automated methodology for a circle target pattern using different values of kernel size and error concentration thresholds.**

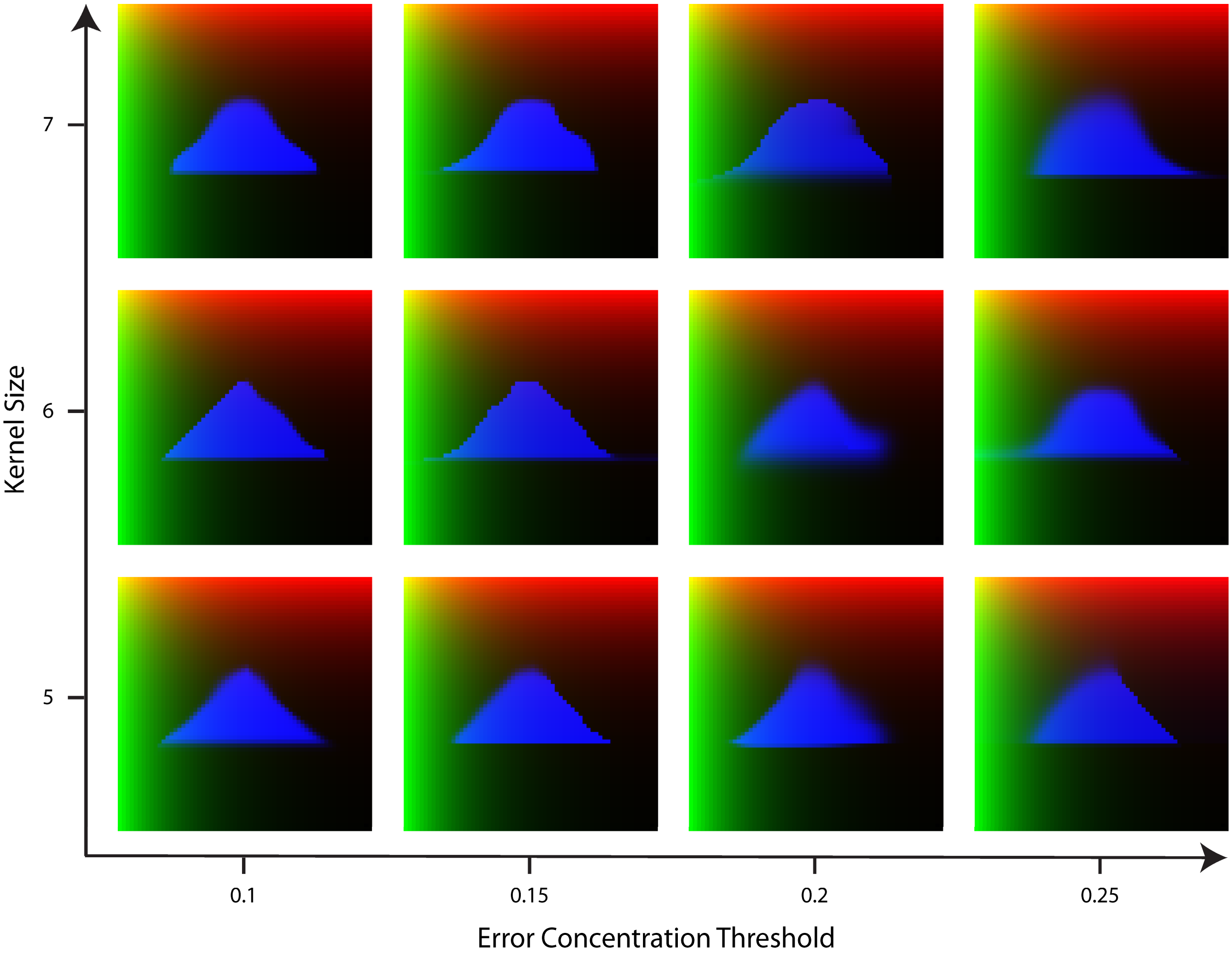


**Supplementary Figure 4. Developed patterns by GRMs discovered with the automated methodology for a triangle target pattern using different values of kernel size and error concentration thresholds.**

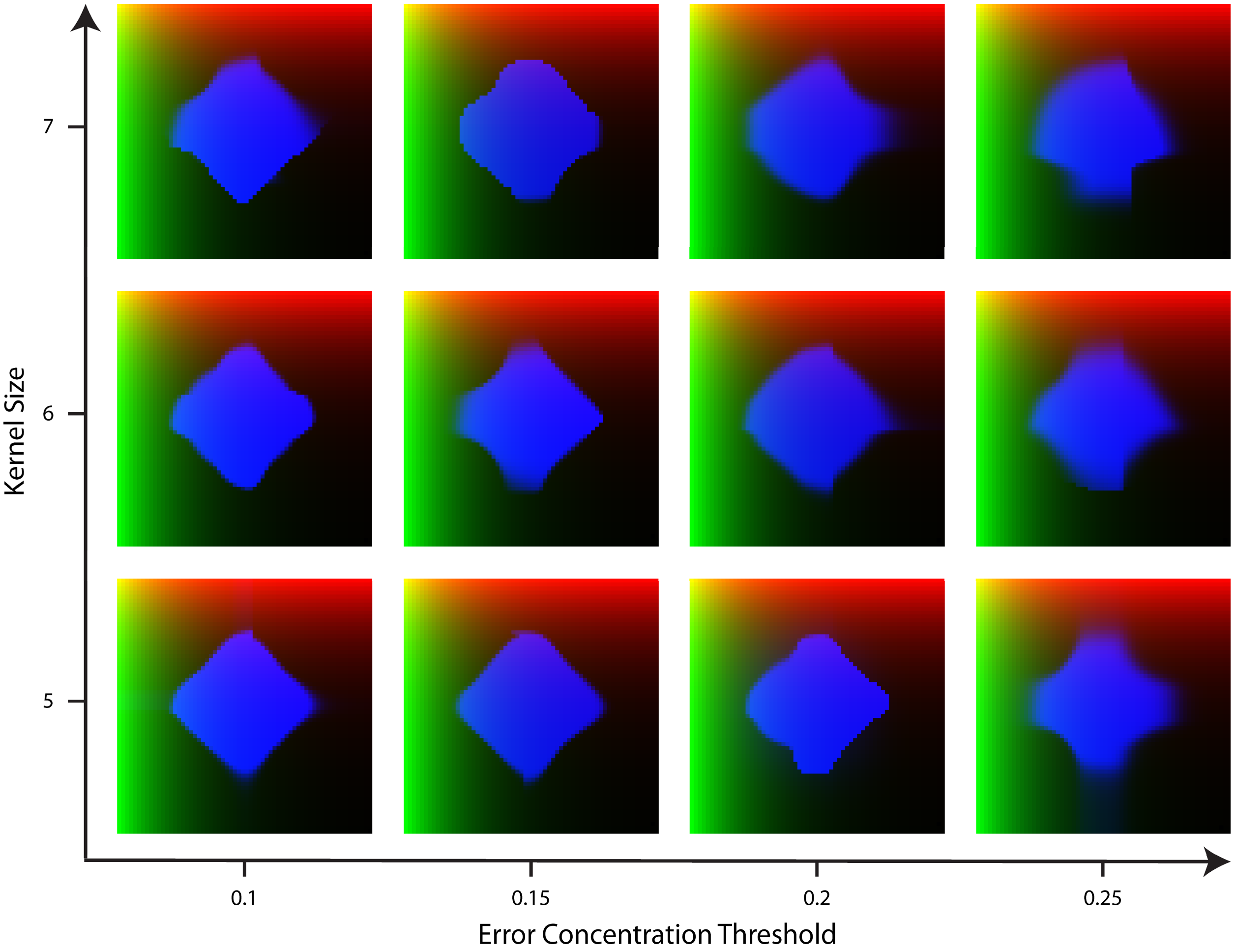


**Supplementary Figure 5. Developed patterns by GRMs discovered with the automated methodology for a diamond target pattern using different values of kernel size and error concentration thresholds.**

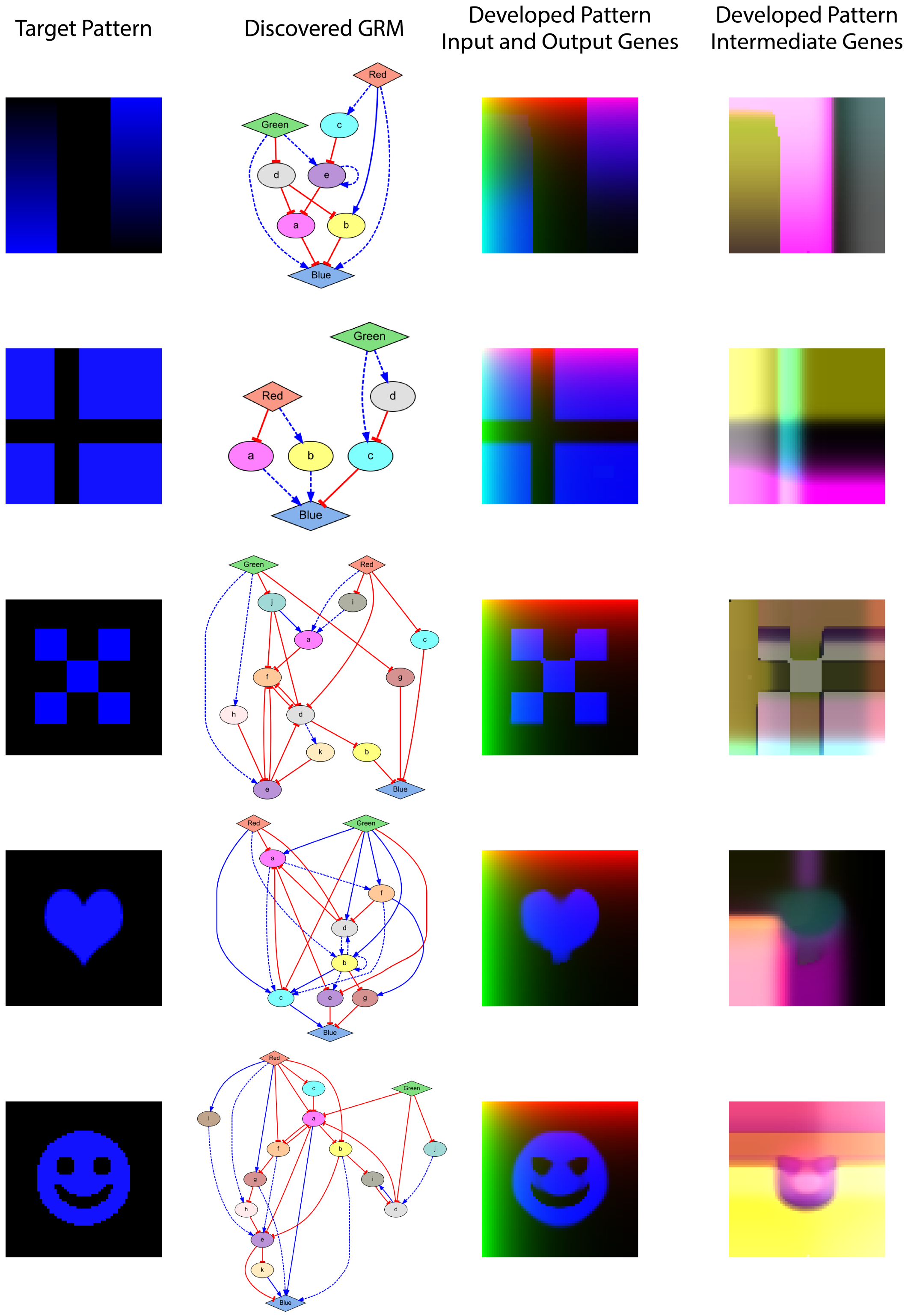


**Supplementary Figure 6. Patterns produced by GRMs discovered by the automated methodology applied to arbitrary shapes, including gradients, flag, checkerboard, and symbols.** All the resulting GRMs correctly produce a steady-state target pattern (blue) with zero error. Expression colors in the developed patterns for the input and output genes and the intermediate genes correspond to the node colors in the GRM network diagrams. Error parameters: $k = 5$, $\alpha = 0.1$, $\beta = 0.001$.

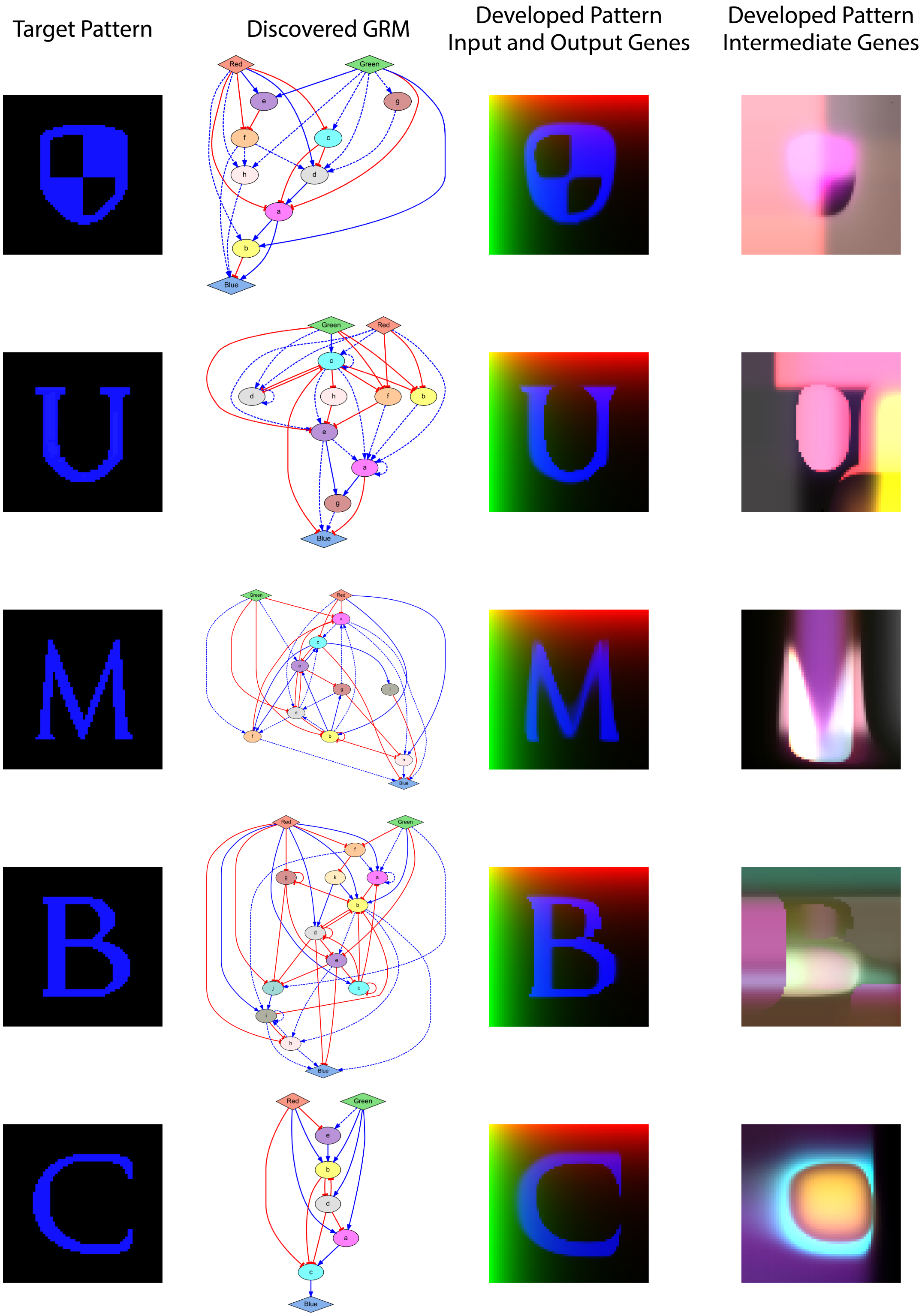


**Supplementary Figure 7. Patterns produced by GRMs discovered by the automated methodology applied to arbitrary shapes, including a logo and characters.** All the resulting GRMs correctly produce a steady-state target pattern (blue) with zero error. Expression colors in the developed patterns for the input and output genes and the intermediate genes correspond to the node colors in the GRM network diagrams. Error parameters: $k = 5$, $\alpha = 0.1$, $\beta = 0.001$.

### 2. System of equations for all GRMs

#### Triangle pattern (Fig. 4B):

$$\frac{\partial Blue}{\partial t} = 0.099 \cdot \frac{90 \cdot e^{6.9} \cdot 62 \cdot d^{9.9}}{(1 + 90 \cdot e^{6.9}) \cdot (1 + 62 \cdot d^{9.9})} - 0.1 \cdot Blue$$

$$\frac{\partial a}{\partial t} = 0.038 \cdot \frac{(3.4 \cdot Green^{4.1} \cdot 28 \cdot c^2 + 3.4 \cdot Green^{4.1} + 28 \cdot c^2)}{(1 + 3.4 \cdot Green^{4.1}) \cdot (1 + 28 \cdot c^2)} - 0.24 \cdot a$$

$$\frac{\partial b}{\partial t} = 0.026 \cdot \frac{\left((5.6 \cdot Red^2 \cdot 2.9 \cdot Green^{2.5} + 5.6 \cdot Red^2 + 2.9 \cdot Green^{2.5}) \cdot 54 \cdot a^{2.8} + (5.6 \cdot Red^2 \cdot 2.9 \cdot Green^{2.5} + 5.6 \cdot Red^2 + 2.9 \cdot Green^{2.5}) + 54 \cdot a^{2.8}\right)}{(1 + 5.6 \cdot Red^2) \cdot (1 + 2.9 \cdot Green^{2.5}) \cdot (1 + 54 \cdot a^{2.8}) \cdot (1 + 17 \cdot d^{2.9}) \cdot (1 + 7.6 \cdot f^{1.5})} - 0.75 \cdot b$$

$$\frac{\partial c}{\partial t} = 0.099 \cdot \frac{1}{(1 + 37 \cdot Green^{3.2}) \cdot (1 + 23 \cdot d^{4.2})} - 0.48 \cdot c$$

$$\frac{\partial d}{\partial t} = 0.053 \cdot \frac{37 \cdot e^{9.6}}{(1 + 37 \cdot e^{9.6}) \cdot (1 + 99 \cdot a^{2.2}) \cdot (1 + 90 \cdot b^{3.2}) \cdot (1 + 62 \cdot c^{1.1})} - 0.43 \cdot d$$

$$\frac{\partial e}{\partial t} = 0.088 \cdot \frac{\left((19 \cdot Red^{9.9} \cdot 42 \cdot b^{6.8} + 19 \cdot Red^{9.9} + 42 \cdot b^{6.8}) \cdot 3.1 \cdot c^{1.6} + (19 \cdot Red^{9.9} \cdot 42 \cdot b^{6.8} + 19 \cdot Red^{9.9} + 42 \cdot b^{6.8}) + 3.1 \cdot c^{1.6}\right)}{(1 + 19 \cdot Red^{9.9}) \cdot (1 + 42 \cdot b^{6.8}) \cdot (1 + 3.1 \cdot c^{1.6})} - 0.93 \cdot e$$

$$\frac{\partial f}{\partial t} = 0.036 \cdot \frac{45 \cdot c^{8.7}}{(1 + 45 \cdot c^{8.7})} - 0.68 \cdot f$$

#### Approximate square pattern (Fig. 5A):

$$\frac{\partial Blue}{\partial t} = 0.096 \cdot \frac{5.8 \cdot a^{8.4}}{(1 + 5.8 \cdot a^{8.4}) \cdot (1 + 2 \cdot Red^{9.6}) \cdot (1 + 2 \cdot Green^{9.6})} - 0.11 \cdot Blue$$

$$\frac{\partial a}{\partial t} = 0.093 \cdot \frac{40 \cdot Red^5 \cdot 40 \cdot Green^5}{(1 + 40 \cdot Red^5) \cdot (1 + 40 \cdot Green^5)} - 0.16 \cdot a$$

#### Approximate circle pattern (Fig. 5B):

$$\frac{\partial Blue}{\partial t} = 0.096 \cdot \frac{51 \cdot a^{7.9}}{(1 + 51 \cdot a^{7.9})} - 0.11 \cdot Blue$$

$$\frac{\partial a}{\partial t} = 0.049 \cdot \frac{45 \cdot b^{5.8}}{(1 + 4.6 \cdot Red^4) \cdot (1 + 4.6 \cdot Green^4) \cdot (1 + 45 \cdot b^{5.8})} - 0.18 \cdot a$$

$$\frac{\partial b}{\partial t} = 0.019 \cdot \frac{36 \cdot Red^3 \cdot 36 \cdot Green^3}{(1 + 36 \cdot Red^3) \cdot (1 + 36 \cdot Green^3)} - 0.65 \cdot b$$

#### Approximate triangle pattern (Fig. 5C):

$$\frac{\partial Blue}{\partial t} = 0.1 \cdot \frac{34 \cdot c^{9.6}}{(1 + 34 \cdot c^{9.6}) \cdot (1 + 94 \cdot a^{9.7})} - 0.11 \cdot Blue$$

$$\frac{\partial a}{\partial t} = 0.073 \cdot \frac{1}{(1 + 36 \cdot Red^9)} - 0.1 \cdot a$$

$$\frac{\partial b}{\partial t} = 0.078 \cdot \frac{17 \cdot Red^{4.4}}{(1 + 17 \cdot Red^{4.4}) \cdot (1 + 45 \cdot Green^{2.5})} - 0.53 \cdot b$$

$$\frac{\partial c}{\partial t} = 0.079 \cdot \frac{1}{(1 + 12 \cdot Red^{1.2}) \cdot (1 + 3.5 \cdot Green^{2.2}) \cdot (1 + 49 \cdot b^{1.9})} - 0.71 \cdot c$$

**Approximate diamond pattern (Fig. 5D):**

$$\frac{\partial Blue}{\partial t} = 0.096 \cdot \frac{44 \cdot a^{9.3} \cdot 33 \cdot Red^{9.4}}{(1 + 44 \cdot a^{9.3}) \cdot (1 + 33 \cdot Red^{9.4})} - 0.1 \cdot Blue$$

$$\frac{\partial a}{\partial t} = 0.013 \cdot \frac{1}{(1 + 3.9 \cdot Red^{2.1}) \cdot (1 + 3.9 \cdot Green^{2.1}) \cdot (1 + 25 \cdot b^{1.6}) \cdot (1 + 51 \cdot c^{4})} - 0.17 \cdot a$$

$$\frac{\partial b}{\partial t} = 0.072 \cdot \frac{28 \cdot d^{7.4}}{(1 + 28 \cdot d^{7.4}) \cdot (1 + 52 \cdot Green^{1.3}) \cdot (1 + 19 \cdot a^{4.3})} - 0.39 \cdot b$$

$$\frac{\partial c}{\partial t} = 0.009 \cdot \frac{1}{(1 + 15 \cdot Red^{9.7}) \cdot (1 + 19 \cdot a^{4.3})} - 0.42 \cdot c$$

$$\frac{\partial d}{\partial t} = 0.07 \cdot \frac{1}{(1 + 14 \cdot Green^{10}) \cdot (1 + 27 \cdot a^{1.7})} - 0.9 \cdot d$$

**Precise square pattern (Fig. 6A):**

$$\frac{\partial Blue}{\partial t} = 0.099 \cdot \frac{1}{(1 + 52 \cdot a^{9.5}) \cdot (1 + 64 \cdot b^{8.1})} - 0.11 \cdot Blue$$

$$\frac{\partial a}{\partial t} = 0.063 \cdot \frac{\left((1.9 \cdot Red^{9.5} \cdot 1.9 \cdot Green^{9.4} + 1.9 \cdot Red^{9.5} + 1.9 \cdot Green^{9.4}) \cdot 30 \cdot c^{5.8} + (1.9 \cdot Red^{9.5} \cdot 1.9 \cdot Green^{9.4} + 1.9 \cdot Red^{9.5} + 1.9 \cdot Green^{9.4}) + 30 \cdot c^{5.8}\right)}{(1 + 1.9 \cdot Red^{9.5}) \cdot (1 + 1.9 \cdot Green^{9.4}) \cdot (1 + 30 \cdot c^{5.8})} - 0.51 \cdot a$$

$$\frac{\partial b}{\partial t} = 0.032 \cdot \frac{1}{(1 + 47 \cdot Red^{9.5})} - 0.63 \cdot b$$

$$\frac{\partial c}{\partial t} = 0.04 \cdot \frac{1}{(1 + 59 \cdot Green^{7.6})} - 0.13 \cdot c$$

**Precise circle pattern (Fig. 6B):**

$$\frac{\partial Blue}{\partial t} = 0.1 \cdot \frac{88 \cdot b^{7.1}}{(1 + 88 \cdot b^{7.1})} - 0.1 \cdot Blue$$

$$\frac{\partial a}{\partial t} = 0.059 \cdot \frac{29 \cdot Green^{2.5} \cdot 29 \cdot Red^{2.5}}{(1 + 29 \cdot Green^{2.5}) \cdot (1 + 29 \cdot Red^{2.5})} - 0.76 \cdot a$$

$$\frac{\partial b}{\partial t} = 0.077 \cdot \frac{16 \cdot a^{4.1}}{(1 + 5.1 \cdot Red^{4.9}) \cdot (1 + 5.1 \cdot Green^{4.9}) \cdot (1 + 16 \cdot a^{4.1}) \cdot (1 + 96 \cdot c^{4.9})} - 0.14 \cdot b$$

$$\frac{\partial c}{\partial t} = 0.0093 \cdot \frac{(9.6 \cdot Red^{4.5} \cdot 9.5 \cdot Green^{4.9} + 9.6 \cdot Red^{4.5} + 9.5 \cdot Green^{4.9})}{(1 + 9.6 \cdot Red^{4.5}) \cdot (1 + 9.5 \cdot Green^{4.9}) \cdot (1 + 14 \cdot a^{6.2})} - 0.99 \cdot c$$

**Precise triangle pattern (Fig. 6C):**

$$\frac{\partial Blue}{\partial t} = 0.1 \cdot \frac{1}{(1 + 80 \cdot a^{9.4}) \cdot (1 + 17 \cdot b^{10}) \cdot (1 + 1\mathrm{e}+02 \cdot c^{10})} - 0.1 \cdot Blue$$

$$\frac{\partial a}{\partial t} = 0.051 \cdot \frac{65 \cdot e^{8.4}}{(1 + 65 \cdot e^{8.4}) \cdot (1 + 69 \cdot Green^{2.4}) \cdot (1 + 95 \cdot b^{1})} - 0.15 \cdot a$$

$$\frac{\partial b}{\partial t} = 0.044 \cdot \frac{1}{(1 + 28 \cdot Red^{3.6}) \cdot (1 + 40 \cdot a^{9.9}) \cdot (1 + 38 \cdot c^{9.9})} - 0.17 \cdot b$$

$$\frac{\partial c}{\partial t} = 0.058 \cdot \frac{\left(\begin{pmatrix}(1.7 \cdot Red^{3.2} \cdot 1.7 \cdot Green^{3.2} + 1.7 \cdot Red^{3.2} + 1.7 \cdot Green^{3.2}) \cdot 17 \cdot a^{1.7} + \\ (1.7 \cdot Red^{3.2} \cdot 1.7 \cdot Green^{3.2} + 1.7 \cdot Red^{3.2} + 1.7 \cdot Green^{3.2}) + 17 \cdot a^{1.7}\end{pmatrix} \cdot 1.7 \cdot d^{3.2} + \begin{pmatrix}(1.7 \cdot Red^{3.2} \cdot 1.7 \cdot Green^{3.2} + 1.7 \cdot Red^{3.2} + 1.7 \cdot Green^{3.2}) \cdot 17 \cdot a^{1.7} + \\ (1.7 \cdot Red^{3.2} \cdot 1.7 \cdot Green^{3.2} + 1.7 \cdot Red^{3.2} + 1.7 \cdot Green^{3.2}) + 17 \cdot a^{1.7}\end{pmatrix} + 1.7 \cdot d^{3.2}\right)}{(1 + 1.7 \cdot Red^{3.2}) \cdot (1 + 1.7 \cdot Green^{3.2}) \cdot (1 + 17 \cdot a^{1.7}) \cdot (1 + 1.7 \cdot d^{3.2}) \cdot (1 + 1\mathrm{e}+02 \cdot b^{1.4})} - 0.2 \cdot c$$

$$\frac{\partial d}{\partial t} = 0.073 \cdot \frac{1}{(1 + 1\mathrm{e}+02 \cdot b^{3.1}) \cdot (1 + 1\mathrm{e}+02 \cdot e^{1})} - 0.53 \cdot d$$

$$\frac{\partial e}{\partial t} = 0.046 \cdot \frac{1}{(1 + 16 \cdot Green^{7.1})} - 0.16 \cdot e$$

**Precise diamond pattern (Fig. 6D):**

$$\frac{\partial Blue}{\partial t} = 0.1 \cdot \frac{(87 \cdot a^{10} \cdot 99 \cdot f^{6.3} + 87 \cdot a^{10} + 99 \cdot f^{6.3})}{(1 + 87 \cdot a^{10}) \cdot (1 + 99 \cdot f^{6.3})} - 0.1 \cdot Blue$$

$$\frac{\partial a}{\partial t} = 0.1 \cdot \frac{1}{(1 + 7.1 \cdot Red^{4.1}) \cdot (1 + 7.1 \cdot Green^{4.1}) \cdot (1 + 98 \cdot b^{2.6}) \cdot (1 + 59 \cdot d^{2.9}) \cdot (1 + 60 \cdot e^{2.4})} - 0.12 \cdot a$$

$$\frac{\partial b}{\partial t} = 0.067 \cdot \frac{1}{(1 + 25 \cdot Red^{2.8}) \cdot (1 + 13 \cdot b^{2.4}) \cdot (1 + 13 \cdot c^{1.5})} - 0.88 \cdot b$$

$$\frac{\partial c}{\partial t} = 0.057 \cdot \frac{1}{(1 + 98 \cdot d^{1.9}) \cdot (1 + 99 \cdot e^{2.8}) \cdot (1 + 98 \cdot g^{7.7})} - 0.48 \cdot c$$

$$\frac{\partial d}{\partial t} = 0.038 \cdot \frac{1}{(1 + 25 \cdot Green^{2.5}) \cdot (1 + 13 \cdot c^{7.9}) \cdot (1 + 11 \cdot d^{5.7})} - 0.38 \cdot d$$

$$\frac{\partial e}{\partial t} = 0.0043 \cdot \frac{(6 \cdot Green^{6.3} \cdot 54 \cdot d^{1.3} + 6 \cdot Green^{6.3} + 54 \cdot d^{1.3}) \cdot 6 \cdot Red^{9.3}}{(1 + 6 \cdot Green^{6.3}) \cdot (1 + 54 \cdot d^{1.3}) \cdot (1 + 6 \cdot Red^{9.3})} - 0.21 \cdot e$$

$$\frac{\partial f}{\partial t} = 0.008 \cdot \frac{(57 \cdot a^{5.4} \cdot 6.1 \cdot c^{6.8} + 57 \cdot a^{5.4} + 6.1 \cdot c^{6.8})}{(1 + 57 \cdot a^{5.4}) \cdot (1 + 6.1 \cdot c^{6.8})} - 0.12 \cdot f$$

$$\frac{\partial g}{\partial t} = 0.092 \cdot \frac{12 \cdot Green^{7.9}}{(1 + 16 \cdot Red^{8.6}) \cdot (1 + 12 \cdot Green^{7.9})} - 0.46 \cdot g$$

**Gradients pattern (Fig. 8A):**

$$\frac{\partial Blue}{\partial t} = 0.095 \cdot \frac{(12 \cdot Red^{1.7} \cdot 5.9 \cdot Green^{9.8} + 12 \cdot Red^{1.7} + 5.9 \cdot Green^{9.8})}{(1 + 12 \cdot Red^{1.7}) \cdot (1 + 5.9 \cdot Green^{9.8}) \cdot (1 + 18 \cdot a^{9.4}) \cdot (1 + 10 \cdot b^{7.5})} - 0.1 \cdot Blue$$

$$\frac{\partial a}{\partial t} = 0.092 \cdot \frac{1}{(1 + 91 \cdot d^{5}) \cdot (1 + 75 \cdot e^{6.4})} - 0.13 \cdot a$$

$$\frac{\partial b}{\partial t} = 0.076 \cdot \frac{39 \cdot Red^{1}}{(1 + 39 \cdot Red^{1}) \cdot (1 + 98 \cdot d^{2.2})} - 0.63 \cdot b$$

$$\frac{\partial c}{\partial t} = 0.025 \cdot \frac{5 \cdot Red^{1.4}}{(1 + 5 \cdot Red^{1.4})} - 0.96 \cdot c$$

$$\frac{\partial d}{\partial t} = 0.087 \cdot \frac{1}{(1 + 31 \cdot Green^{3.7})} - 0.98 \cdot d$$

$$\frac{\partial e}{\partial t} = 0.048 \cdot \frac{(2.6 \cdot Green^{4.1} \cdot 95 \cdot e^{5} + 2.6 \cdot Green^{4.1} + 95 \cdot e^{5})}{(1 + 2.6 \cdot Green^{4.1}) \cdot (1 + 95 \cdot e^{5}) \cdot (1 + 49 \cdot c^{7.6})} - 0.99 \cdot e$$

**Flag pattern (Fig. 8A):**

$$\frac{\partial Blue}{\partial t} = 0.098 \cdot \frac{(90 \cdot a^{9.9} \cdot 18 \cdot b^{8.1} + 90 \cdot a^{9.9} + 18 \cdot b^{8.1})}{(1 + 90 \cdot a^{9.9}) \cdot (1 + 18 \cdot b^{8.1}) \cdot (1 + 16 \cdot c^{9.7})} - 0.1 \cdot Blue$$

$$\frac{\partial a}{\partial t} = 0.076 \cdot \frac{1}{(1 + 48 \cdot Red^{3.1})} - 0.22 \cdot a$$

$$\frac{\partial b}{\partial t} = 0.041 \cdot \frac{8.1 \cdot Red^{9.7}}{(1 + 8.1 \cdot Red^{9.7})} - 0.36 \cdot b$$

$$\frac{\partial c}{\partial t} = 0.046 \cdot \frac{8.1 \cdot Green^{9.9}}{(1 + 8.1 \cdot Green^{9.9}) \cdot (1 + 16 \cdot d^{9.7})} - 0.33 \cdot c$$

$$\frac{\partial d}{\partial t} = 0.061 \cdot \frac{3.5 \cdot Green^{3.1}}{(1 + 3.5 \cdot Green^{3.1})} - 0.34 \cdot d$$

### Checkerboard pattern (Fig. 8A):

$$\frac{\partial Blue}{\partial t} = 0.1 \cdot \frac{1}{(1 + 74 \cdot b^{7}) \cdot (1 + 74 \cdot c^{4.9}) \cdot (1 + 73 \cdot g^{6})} - 0.1 \cdot Blue$$

$$\frac{\partial a}{\partial t} = 0.053 \cdot \frac{(3.4 \cdot Red^{5.1} \cdot 80 \cdot i^{4.9} + 3.4 \cdot Red^{5.1} + 80 \cdot i^{4.9}) \cdot 54 \cdot j^{9.3}}{(1 + 3.4 \cdot Red^{5.1}) \cdot (1 + 80 \cdot i^{4.9}) \cdot (1 + 54 \cdot j^{9.3})} - 0.25 \cdot a$$

$$\frac{\partial b}{\partial t} = 0.084 \cdot \frac{1}{(1 + 76 \cdot d^{6.6})} - 0.21 \cdot b$$

$$\frac{\partial c}{\partial t} = 0.089 \cdot \frac{1}{(1 + 67 \cdot Red^{7.3})} - 0.12 \cdot c$$

$$\frac{\partial d}{\partial t} = 0.09 \cdot \frac{1}{(1 + 3.2 \cdot Red^{9.9}) \cdot (1 + 89 \cdot e^{9.1}) \cdot (1 + 13 \cdot f^{8.3}) \cdot (1 + 4 \cdot j^{3.6})} - 0.17 \cdot d$$

$$\frac{\partial e}{\partial t} = 0.054 \cdot \frac{12 \cdot Green^{5.4}}{(1 + 12 \cdot Green^{5.4}) \cdot (1 + 22 \cdot f^{8.7}) \cdot (1 + 94 \cdot h^{8.5}) \cdot (1 + 93 \cdot k^{8.4})} - 0.95 \cdot e$$

$$\frac{\partial f}{\partial t} = 0.095 \cdot \frac{1}{(1 + 67 \cdot a^{8.3}) \cdot (1 + 4 \cdot d^{4.1}) \cdot (1 + 87 \cdot e^{9.5}) \cdot (1 + 4 \cdot j^{3.7})} - 0.73 \cdot f$$

$$\frac{\partial g}{\partial t} = 0.08 \cdot \frac{1}{(1 + 59 \cdot Green^{9})} - 0.15 \cdot g$$

$$\frac{\partial h}{\partial t} = 0.086 \cdot \frac{2.5 \cdot Green^{3.6}}{(1 + 2.5 \cdot Green^{3.6})} - 0.29 \cdot h$$

$$\frac{\partial i}{\partial t} = 0.078 \cdot \frac{1}{(1 + 51 \cdot Red^{2.8})} - 0.36 \cdot i$$

$$\frac{\partial j}{\partial t} = 0.097 \cdot \frac{1}{(1 + 2.8 \cdot Green^{9.2})} - 0.75 \cdot j$$

$$\frac{\partial k}{\partial t} = 0.093 \cdot \frac{3 \cdot d^{8.2}}{(1 + 3 \cdot d^{8.2})} - 0.32 \cdot k$$

### Heart pattern (Fig. 8A):

$$\frac{\partial Blue}{\partial t} = 0.1 \cdot \frac{18 \cdot c^{10}}{(1 + 18 \cdot c^{10}) \cdot (1 + 23 \cdot e^{7.4}) \cdot (1 + 9 \cdot g^{2.6})} - 0.1 \cdot Blue$$

$$\frac{\partial a}{\partial t} = 0.084 \cdot \frac{29 \cdot Green^{5.2}}{(1 + 9 \cdot Red^{2.6}) \cdot (1 + 29 \cdot Green^{5.2}) \cdot (1 + 6.6 \cdot c^{9.5}) \cdot (1 + 63 \cdot d^{1.5})} - 0.23 \cdot a$$

$$\frac{\partial b}{\partial t} = 0.031 \cdot \frac{\left((13 \cdot Red^{1.8} \cdot 19 \cdot b^{9.9} + 13 \cdot Red^{1.8} + 19 \cdot b^{9.9}) \cdot 64 \cdot d^{6.4} + (13 \cdot Red^{1.8} \cdot 19 \cdot b^{9.9} + 13 \cdot Red^{1.8} + 19 \cdot b^{9.9}) + 64 \cdot d^{6.4}\right) \cdot 23 \cdot Green^{2.8}}{(1 + 13 \cdot Red^{1.8}) \cdot (1 + 19 \cdot b^{9.9}) \cdot (1 + 64 \cdot d^{6.4}) \cdot (1 + 23 \cdot Green^{2.8})} - 0.46 \cdot b$$

$$\frac{\partial c}{\partial t} = 0.055 \cdot \frac{(36 \cdot a^{3.3} \cdot 19 \cdot f^{9.9} + 36 \cdot a^{3.3} + 19 \cdot f^{9.9}) \cdot 18 \cdot Red^{2.5} \cdot 49 \cdot b^{3.1}}{(1 + 36 \cdot a^{3.3}) \cdot (1 + 19 \cdot f^{9.9}) \cdot (1 + 18 \cdot Red^{2.5}) \cdot (1 + 4.5 \cdot Green^{3.2}) \cdot (1 + 49 \cdot b^{3.1})} - 0.2 \cdot c$$

$$\frac{\partial d}{\partial t} = 0.027 \cdot \frac{79 \cdot b^{7.9} \cdot 18 \cdot Green^{8.3}}{(1 + 79 \cdot b^{7.9}) \cdot (1 + 23 \cdot Red^{1.5}) \cdot (1 + 18 \cdot Green^{8.3}) \cdot (1 + 85 \cdot f^{3.9})} - 0.95 \cdot d$$

$$\frac{\partial e}{\partial t} = 0.083 \cdot \frac{18 \cdot b^{10}}{(1 + 18 \cdot b^{10}) \cdot (1 + 7.4 \cdot Green^{6.9}) \cdot (1 + 55 \cdot a^{3.7})} - 0.13 \cdot e$$

$$\frac{\partial f}{\partial t} = 0.081 \cdot \frac{7.9 \cdot a^{10} \cdot 4.1 \cdot Green^{7.5}}{(1 + 7.9 \cdot a^{10}) \cdot (1 + 4.1 \cdot Green^{7.5})} - 0.11 \cdot f$$

$$\frac{\partial g}{\partial t} = 0.088 \cdot \frac{18 \cdot f^{2.4}}{(1 + 35 \cdot b^{9.8}) \cdot (1 + 18 \cdot f^{2.4})} - 0.35 \cdot g$$

**Smiley face pattern (Fig. 8A):**

$$\frac{\partial Blue}{\partial t} = 0.1 \cdot \frac{(49 \cdot b^{9.5} \cdot 99 \cdot g^{6.5} + 49 \cdot b^{9.5} + 99 \cdot g^{6.5}) \cdot 86 \cdot a^{10} \cdot 53 \cdot k^{8.5}}{(1 + 49 \cdot b^{9.5}) \cdot (1 + 99 \cdot g^{6.5}) \cdot (1 + 86 \cdot a^{10}) \cdot (1 + 5.2 \cdot e^{9.9}) \cdot (1 + 53 \cdot k^{8.5})} - 0.1 \cdot Blue$$

$$\frac{\partial a}{\partial t} = 0.066 \cdot \frac{1}{(1 + 5.2 \cdot Red^{3.6}) \cdot (1 + 5.2 \cdot Green^{3.9}) \cdot (1 + 84 \cdot c^{2.3}) \cdot (1 + 59 \cdot d^{2.9}) \cdot (1 + 5.2 \cdot f^{3.1})} - 0.22 \cdot a$$

$$\frac{\partial b}{\partial t} = 0.098 \cdot \frac{1}{(1 + 7.3 \cdot Red^{9.5}) \cdot (1 + 23 \cdot a^{6.2})} - 0.18 \cdot b$$

$$\frac{\partial c}{\partial t} = 0.044 \cdot \frac{1}{(1 + 41 \cdot Red^{2.4})} - 0.47 \cdot c$$

$$\frac{\partial d}{\partial t} = 0.062 \cdot \frac{71 \cdot j^{5.7}}{(1 + 71 \cdot j^{5.7}) \cdot (1 + 70 \cdot Green^{1.7}) \cdot (1 + 6.1 \cdot i^{5.4})} - 0.38 \cdot d$$

$$\frac{\partial e}{\partial t} = 0.094 \cdot \frac{(53 \cdot f^{9.5} \cdot 69 \cdot l^{9.2} + 53 \cdot f^{9.5} + 69 \cdot l^{9.2})}{(1 + 53 \cdot f^{9.5}) \cdot (1 + 69 \cdot l^{9.2}) \cdot (1 + 5.2 \cdot a^{9.3}) \cdot (1 + 81 \cdot b^{1.7}) \cdot (1 + 5.2 \cdot h^{4.5})} - 0.31 \cdot e$$

$$\frac{\partial f}{\partial t} = 0.077 \cdot \frac{1}{(1 + 6.1 \cdot Red^{5.4}) \cdot (1 + 6.1 \cdot a^{3.1})} - 0.98 \cdot f$$

$$\frac{\partial g}{\partial t} = 0.076 \cdot \frac{7.3 \cdot Red^{4.4}}{(1 + 7.3 \cdot Red^{4.4}) \cdot (1 + 18 \cdot f^{8})} - 0.16 \cdot g$$

$$\frac{\partial h}{\partial t} = 0.062 \cdot \frac{16 \cdot Red^{9.9}}{(1 + 16 \cdot Red^{9.9}) \cdot (1 + 5.9 \cdot g^{7.8})} - 0.33 \cdot h$$

$$\frac{\partial i}{\partial t} = 0.025 \cdot \frac{85 \cdot d^{9}}{(1 + 5.2 \cdot b^{3.1}) \cdot (1 + 85 \cdot d^{9})} - 0.32 \cdot i$$

$$\frac{\partial j}{\partial t} = 0.074 \cdot \frac{1}{(1 + 14 \cdot Green^{4})} - 0.56 \cdot j$$

$$\frac{\partial k}{\partial t} = 0.078 \cdot \frac{1}{(1 + 9 \cdot e^{2.9})} - 0.37 \cdot k$$

$$\frac{\partial l}{\partial t} = 0.075 \cdot \frac{1.7 \cdot Red^{7}}{(1 + 1.7 \cdot Red^{7})} - 0.43 \cdot l$$

**Logo pattern (Fig. 8B):**

$$\frac{\partial Blue}{\partial t} = 0.1 \cdot \frac{(58 \cdot f^{8.8} \cdot 9.7 \cdot h^{10} + 58 \cdot f^{8.8} + 9.7 \cdot h^{10}) \cdot 79 \cdot a^{6.8}}{(1 + 58 \cdot f^{8.8}) \cdot (1 + 9.7 \cdot h^{10}) \cdot (1 + 79 \cdot a^{6.8}) \cdot (1 + 97 \cdot b^{9.1})} - 0.1 \cdot Blue$$

$$\frac{\partial a}{\partial t} = 0.083 \cdot \frac{16 \cdot d^{6.4}}{(1 + 3.6 \cdot Red^{6.7}) \cdot (1 + 3.6 \cdot Green^{8.1}) \cdot (1 + 47 \cdot c^{2.3}) \cdot (1 + 16 \cdot d^{6.4})} - 0.18 \cdot a$$

$$\frac{\partial b}{\partial t} = 0.019 \cdot \frac{11 \cdot Red^{9.5} \cdot 10 \cdot Green^{9.9} \cdot 13 \cdot a^{10}}{(1 + 11 \cdot Red^{9.5}) \cdot (1 + 10 \cdot Green^{9.9}) \cdot (1 + 13 \cdot a^{10})} - 0.84 \cdot b$$

$$\frac{\partial c}{\partial t} = 0.004 \cdot \frac{6.8 \cdot Green^{6.1}}{(1 + 6.8 \cdot Green^{6.1}) \cdot (1 + 1.7 \cdot Red^{6.1})} - 0.59 \cdot c$$

$$\frac{\partial d}{\partial t} = 0.022 \cdot \frac{\left((38 \cdot Green^{2.8} \cdot 14 \cdot f^{7.7} + 38 \cdot Green^{2.8} + 14 \cdot f^{7.7}) \cdot 39 \cdot g^{7.5} + (38 \cdot Green^{2.8} \cdot 14 \cdot f^{7.7} + 38 \cdot Green^{2.8} + 14 \cdot f^{7.7}) + 39 \cdot g^{7.5}\right) \cdot 36 \cdot Red^{1.4}}{(1 + 38 \cdot Green^{2.8}) \cdot (1 + 14 \cdot f^{7.7}) \cdot (1 + 39 \cdot g^{7.5}) \cdot (1 + 36 \cdot Red^{1.4}) \cdot (1 + 97 \cdot c^{3})} - 0.29 \cdot d$$

$$\frac{\partial e}{\partial t} = 0.03 \cdot \frac{22 \cdot Red^{1.9} \cdot 24 \cdot Green^{4.5}}{(1 + 22 \cdot Red^{1.9}) \cdot (1 + 24 \cdot Green^{4.5})} - 0.94 \cdot e$$

$$\frac{\partial f}{\partial t} = 0.03 \cdot \frac{1}{(1 + 3.6 \cdot Red^{6.7}) \cdot (1 + 92 \cdot e^{9.1})} - 0.84 \cdot f$$

$$\frac{\partial g}{\partial t} = 0.091 \cdot \frac{10 \cdot Green^{9.9}}{(1 + 10 \cdot Green^{9.9})} - 0.22 \cdot g$$

$$\frac{\partial h}{\partial t} = 0.076 \cdot \frac{\left((11 \cdot Red^{10} \cdot 9.6 \cdot Green^{9.9} + 11 \cdot Red^{10} + 9.6 \cdot Green^{9.9}) \cdot 39 \cdot f^{10} + (11 \cdot Red^{10} \cdot 9.6 \cdot Green^{9.9} + 11 \cdot Red^{10} + 9.6 \cdot Green^{9.9}) + 39 \cdot f^{10}\right)}{(1 + 11 \cdot Red^{10}) \cdot (1 + 9.6 \cdot Green^{9.9}) \cdot (1 + 39 \cdot f^{10})} - 0.4 \cdot h$$

### Letter ‘U’ pattern (Fig. 8B):

$$\frac{\partial Blue}{\partial t} = 0.1 \cdot \frac{(49 \cdot e^{9.9} \cdot 72 \cdot g^{4.4} + 49 \cdot e^{9.9} + 72 \cdot g^{4.4})}{(1 + 49 \cdot e^{9.9}) \cdot (1 + 72 \cdot g^{4.4}) \cdot (1 + 1\mathrm{e}+02 \cdot a^{9.8}) \cdot (1 + 80 \cdot c^{10})} - 0.1 \cdot Blue$$

$$\frac{\partial a}{\partial t} = 0.0027 \cdot \frac{\left( \left( \left( \begin{matrix} ((2.6 \cdot Red^{9.9} \cdot 57 \cdot a^{5.3} + 2.6 \cdot Red^{9.9} + 57 \cdot a^{5.3}) \cdot 95 \cdot b^{9.3} + (2.6 \cdot Red^{9.9} \cdot 57 \cdot a^{5.3} + 2.6 \cdot Red^{9.9} + 57 \cdot a^{5.3}) + 95 \cdot b^{9.3}) \cdot 93 \cdot c^{9.8} \\ +((2.6 \cdot Red^{9.9} \cdot 57 \cdot a^{5.3} + 2.6 \cdot Red^{9.9} + 57 \cdot a^{5.3}) \cdot 95 \cdot b^{9.3} + (2.6 \cdot Red^{9.9} \cdot 57 \cdot a^{5.3} + 2.6 \cdot Red^{9.9} + 57 \cdot a^{5.3}) + 95 \cdot b^{9.3}) + 93 \cdot c^{9.8} \end{matrix} \right) \cdot 5.2 \cdot e^{6.3} + \left( \begin{matrix} ((2.6 \cdot Red^{9.9} \cdot 57 \cdot a^{5.3} + 2.6 \cdot Red^{9.9} + 57 \cdot a^{5.3}) \cdot 95 \cdot b^{9.3} + (2.6 \cdot Red^{9.9} \cdot 57 \cdot a^{5.3} + 2.6 \cdot Red^{9.9} + 57 \cdot a^{5.3}) + 95 \cdot b^{9.3}) \cdot 93 \cdot c^{9.8} \\ +((2.6 \cdot Red^{9.9} \cdot 57 \cdot a^{5.3} + 2.6 \cdot Red^{9.9} + 57 \cdot a^{5.3}) \cdot 95 \cdot b^{9.3} + (2.6 \cdot Red^{9.9} \cdot 57 \cdot a^{5.3} + 2.6 \cdot Red^{9.9} + 57 \cdot a^{5.3}) + 95 \cdot b^{9.3}) + 93 \cdot c^{9.8} \end{matrix} \right) + 5.2 \cdot e^{6.3} \right) \cdot 95 \cdot f^{10} + \left( \left( \begin{matrix} ((2.6 \cdot Red^{9.9} \cdot 57 \cdot a^{5.3} + 2.6 \cdot Red^{9.9} + 57 \cdot a^{5.3}) \cdot 95 \cdot b^{9.3} + (2.6 \cdot Red^{9.9} \cdot 57 \cdot a^{5.3} + 2.6 \cdot Red^{9.9} + 57 \cdot a^{5.3}) + 95 \cdot b^{9.3}) \cdot 93 \cdot c^{9.8} \\ +((2.6 \cdot Red^{9.9} \cdot 57 \cdot a^{5.3} + 2.6 \cdot Red^{9.9} + 57 \cdot a^{5.3}) \cdot 95 \cdot b^{9.3} + (2.6 \cdot Red^{9.9} \cdot 57 \cdot a^{5.3} + 2.6 \cdot Red^{9.9} + 57 \cdot a^{5.3}) + 95 \cdot b^{9.3}) + 93 \cdot c^{9.8} \end{matrix} \right) \cdot 5.2 \cdot e^{6.3} + \left( \begin{matrix} \left( \begin{matrix} (2.6 \cdot Red^{9.9} \cdot 57 \cdot a^{5.3} + 2.6 \cdot Red^{9.9} + 57 \cdot a^{5.3}) \cdot 95 \cdot b^{9.3} + (2.6 \cdot Red^{9.9} \cdot 57 \cdot a^{5.3} + 2.6 \cdot Red^{9.9} + 57 \cdot a^{5.3}) + \\ 95 \cdot b^{9.3} \end{matrix} \right) \cdot 93 \cdot c^{9.8} \\ +((2.6 \cdot Red^{9.9} \cdot 57 \cdot a^{5.3} + 2.6 \cdot Red^{9.9} + 57 \cdot a^{5.3}) \cdot 95 \cdot b^{9.3} + (2.6 \cdot Red^{9.9} \cdot 57 \cdot a^{5.3} + 2.6 \cdot Red^{9.9} + 57 \cdot a^{5.3}) + 95 \cdot b^{9.3}) + 93 \cdot c^{9.8} \end{matrix} \right) + 5.2 \cdot e^{6.3} \right) + 95 \cdot f^{10} \right)}{(1 + 2.6 \cdot Red^{9.9}) \cdot (1 + 57 \cdot a^{5.3}) \cdot (1 + 95 \cdot b^{9.3}) \cdot (1 + 93 \cdot c^{9.8}) \cdot (1 + 5.2 \cdot e^{6.3}) \cdot (1 + 95 \cdot f^{10})} - 0.15 \cdot a$$

$$\frac{\partial b}{\partial t} = 0.096 \cdot \frac{1}{(1 + 3.6 \cdot Red^{5.3}) \cdot (1 + 53 \cdot Green^{6.4}) \cdot (1 + 96 \cdot c^{1.3})} - 0.22 \cdot b$$

$$\frac{\partial c}{\partial t} = 0.078 \cdot \frac{(20 \cdot Red^{5} \cdot 30 \cdot c^{3.6} + 20 \cdot Red^{5} + 30 \cdot c^{3.6}) \cdot 17 \cdot Green^{7.2}}{(1 + 20 \cdot Red^{5}) \cdot (1 + 30 \cdot c^{3.6}) \cdot (1 + 17 \cdot Green^{7.2}) \cdot (1 + 77 \cdot d^{1.6})} - 0.78 \cdot c$$

$$\frac{\partial d}{\partial t} = 0.084 \cdot \frac{((2.7 \cdot Red^{5.4} \cdot 5.2 \cdot Green^{6.8} + 2.7 \cdot Red^{5.4} + 5.2 \cdot Green^{6.8}) \cdot 9.8 \cdot d^{5.6} + (2.7 \cdot Red^{5.4} \cdot 5.2 \cdot Green^{6.8} + 2.7 \cdot Red^{5.4} + 5.2 \cdot Green^{6.8}) + 9.8 \cdot d^{5.6})}{(1 + 2.7 \cdot Red^{5.4}) \cdot (1 + 5.2 \cdot Green^{6.8}) \cdot (1 + 9.8 \cdot d^{5.6}) \cdot (1 + 43 \cdot c^{1.9})} - 0.85 \cdot d$$

$$\frac{\partial e}{\partial t} = 0.033 \cdot \frac{(29 \cdot Red^{4.4} \cdot 94 \cdot c^{1.7} + 29 \cdot Red^{4.4} + 94 \cdot c^{1.7})}{(1 + 29 \cdot Red^{4.4}) \cdot (1 + 94 \cdot c^{1.7}) \cdot (1 + 4.8 \cdot Green^{3.4}) \cdot (1 + 85 \cdot f^{1.6}) \cdot (1 + 37 \cdot h^{8.6})} - 0.27 \cdot e$$

$$\frac{\partial f}{\partial t} = 0.091 \cdot \frac{1}{(1 + 70 \cdot Red^{2.7}) \cdot (1 + 19 \cdot Green^{5.7}) \cdot (1 + 97 \cdot c^{1.3})} - 0.49 \cdot f$$

$$\frac{\partial g}{\partial t} = 0.096 \cdot \frac{89 \cdot a^{2.7} \cdot 72 \cdot e^{9.3}}{(1 + 89 \cdot a^{2.7}) \cdot (1 + 72 \cdot e^{9.3})} - 0.1 \cdot g$$

$$\frac{\partial h}{\partial t} = 0.01 \cdot \frac{1}{(1 + 19 \cdot c^{1})} - 0.57 \cdot h$$

### Letter ‘M’ pattern (Fig. 8B):

$$\frac{\partial Blue}{\partial t} = 0.098 \cdot \frac{(48 \cdot a^{10} \cdot 50 \cdot f^{9.9} + 48 \cdot a^{10} + 50 \cdot f^{9.9}) \cdot 43 \cdot h^{9.3}}{(1 + 48 \cdot a^{10}) \cdot (1 + 50 \cdot f^{9.9}) \cdot (1 + 11 \cdot c^{10}) \cdot (1 + 43 \cdot h^{9.3}) \cdot (1 + 11 \cdot i^{6.3})} - 0.11 \cdot Blue$$

$$\frac{\partial a}{\partial t} = 0.039 \cdot \frac{(61 \cdot b^{4.6} \cdot 85 \cdot g^{9.8} + 61 \cdot b^{4.6} + 85 \cdot g^{9.8})}{(1 + 61 \cdot b^{4.6}) \cdot (1 + 85 \cdot g^{9.8}) \cdot (1 + 4.3 \cdot Red^{1.1}) \cdot (1 + 1.7 \cdot Green^{7.8}) \cdot (1 + 3.8 \cdot e^{2.2}) \cdot (1 + 1.7 \cdot f^{5.7})} - 0.69 \cdot a$$

$$\frac{\partial b}{\partial t} = 0.044 \cdot \frac{86 \cdot Red^{5.4}}{(1 + 86 \cdot Red^{5.4}) \cdot (1 + 3.8 \cdot Green^{2.1}) \cdot (1 + 12 \cdot b^{10}) \cdot (1 + 83 \cdot d^{1.9})} - 0.99 \cdot b$$

$$\frac{\partial c}{\partial t} = 0.091 \cdot \frac{(38 \cdot a^{8.8} \cdot 98 \cdot d^{1} + 38 \cdot a^{8.8} + 98 \cdot d^{1}) \cdot 49 \cdot e^{9.9}}{(1 + 38 \cdot a^{8.8}) \cdot (1 + 98 \cdot d^{1}) \cdot (1 + 4.7 \cdot Red^{1.3}) \cdot (1 + 49 \cdot e^{9.9})} - 0.66 \cdot c$$

$$\frac{\partial d}{\partial t} = 0.085 \cdot \frac{((43 \cdot Red^{2.3} \cdot 46 \cdot b^{6.3} + 43 \cdot Red^{2.3} + 46 \cdot b^{6.3}) \cdot 18 \cdot g^{6.1} + (43 \cdot Red^{2.3} \cdot 46 \cdot b^{6.3} + 43 \cdot Red^{2.3} + 46 \cdot b^{6.3}) + 18 \cdot g^{6.1})}{(1 + 43 \cdot Red^{2.3}) \cdot (1 + 46 \cdot b^{6.3}) \cdot (1 + 18 \cdot g^{6.1}) \cdot (1 + 83 \cdot Green^{2.7}) \cdot (1 + 9.1 \cdot e^{9.8})} - 0.44 \cdot d$$

$$\frac{\partial e}{\partial t} = 0.096 \cdot \frac{18 \cdot Green^{6.1} \cdot 34 \cdot b^{10}}{(1 + 18 \cdot Green^{6.1}) \cdot (1 + 34 \cdot b^{10}) \cdot (1 + 3.8 \cdot c^{1.3}) \cdot (1 + 75 \cdot d^{2.5})} - 0.3 \cdot e$$

$$\frac{\partial f}{\partial t} = 0.056 \cdot \frac{(7.6 \cdot Green^{7.9} \cdot 79 \cdot d^{3.5} + 7.6 \cdot Green^{7.9} + 79 \cdot d^{3.5}) \cdot 62 \cdot c^{2} \cdot 98 \cdot e^{2}}{(1 + 7.6 \cdot Green^{7.9}) \cdot (1 + 79 \cdot d^{3.5}) \cdot (1 + 62 \cdot c^{2}) \cdot (1 + 98 \cdot e^{2})} - 0.16 \cdot f$$

$$\frac{\partial g}{\partial t} = 0.017 \cdot \frac{43 \cdot b^{8.8}}{(1 + 43 \cdot b^{8.8}) \cdot (1 + 4.3 \cdot e^{6.9})} - 0.27 \cdot g$$

$$\frac{\partial h}{\partial t} = 0.076 \cdot \frac{44 \cdot a^{9.9} \cdot 34 \cdot Red^{7.9}}{(1 + 44 \cdot a^{9.9}) \cdot (1 + 34 \cdot Red^{7.9}) \cdot (1 + 23 \cdot b^{8.8}) \cdot (1 + 9.8 \cdot g^{3.1})} - 0.34 \cdot h$$

$$\frac{\partial i}{\partial t} = 0.029 \cdot \frac{8.5 \cdot c^{5.3}}{(1 + 8.5 \cdot c^{5.3})} - 0.13 \cdot i$$

### Letter ‘B’ pattern (Fig. 8B):

$$\frac{\partial Blue}{\partial t} = 0.1 \cdot \frac{\left((50 \cdot b^{9.6} \cdot 90 \cdot h^{9.5} + 50 \cdot b^{9.6} + 90 \cdot h^{9.5}) \cdot 50 \cdot i^{9.2} + (50 \cdot b^{9.6} \cdot 90 \cdot h^{9.5} + 50 \cdot b^{9.6} + 90 \cdot h^{9.5}) + 50 \cdot i^{9.2}\right)}{(1 + 50 \cdot b^{9.6}) \cdot (1 + 90 \cdot h^{9.5}) \cdot (1 + 50 \cdot i^{9.2}) \cdot (1 + 18 \cdot d^{10}) \cdot (1 + 15 \cdot e^{8.7})} - 0.1 \cdot Blue$$

$$\frac{\partial a}{\partial t} = 0.093 \cdot \frac{\left((1.5 \cdot Green^{5.4} \cdot 78 \cdot a^{3.5} + 1.5 \cdot Green^{5.4} + 78 \cdot a^{3.5}) \cdot 38 \cdot f^{9.8} + (1.5 \cdot Green^{5.4} \cdot 78 \cdot a^{3.5} + 1.5 \cdot Green^{5.4} + 78 \cdot a^{3.5}) + 38 \cdot f^{9.8}\right) \cdot 44 \cdot Red^{5.9}}{(1 + 1.5 \cdot Green^{5.4}) \cdot (1 + 78 \cdot a^{3.5}) \cdot (1 + 38 \cdot f^{9.8}) \cdot (1 + 44 \cdot Red^{5.9}) \cdot (1 + 86 \cdot c^{1.2})} - 0.53 \cdot a$$

$$\frac{\partial b}{\partial t} = 0.066 \cdot \frac{38 \cdot Red^{9.9} \cdot 19 \cdot Green^{2} \cdot 38 \cdot k^{8.2}}{(1 + 38 \cdot Red^{9.9}) \cdot (1 + 19 \cdot Green^{2}) \cdot (1 + 67 \cdot a^{1}) \cdot (1 + 96 \cdot c^{1.2}) \cdot (1 + 15 \cdot d^{9.2}) \cdot (1 + 11 \cdot g^{7.8}) \cdot (1 + 6.6 \cdot i^{1.1}) \cdot (1 + 38 \cdot k^{8.2})} - 0.72 \cdot b$$

$$\frac{\partial c}{\partial t} = 0.083 \cdot \frac{1.5 \cdot Red^{7.2}}{(1 + 1.5 \cdot Red^{7.2}) \cdot (1 + 16 \cdot c^{1.8}) \cdot (1 + 22 \cdot e^{9.5})} - 0.14 \cdot c$$

$$\frac{\partial d}{\partial t} = 0.083 \cdot \frac{15 \cdot Red^{7.2} \cdot 81 \cdot k^{7.3}}{(1 + 15 \cdot Red^{7.2}) \cdot (1 + 94 \cdot b^{1.7}) \cdot (1 + 38 \cdot c^{8.1}) \cdot (1 + 6.6 \cdot d^{3.1}) \cdot (1 + 81 \cdot k^{7.3})} - 0.43 \cdot d$$

$$\frac{\partial e}{\partial t} = 0.07 \cdot \frac{24 \cdot b^{6.9}}{(1 + 24 \cdot b^{6.9}) \cdot (1 + 6.6 \cdot Green^{6.1}) \cdot (1 + 1\mathrm{e}{+}02 \cdot d^{9.6}) \cdot (1 + 68 \cdot g^{9.2})} - 0.38 \cdot e$$

$$\frac{\partial f}{\partial t} = 0.025 \cdot \frac{1}{(1 + 85 \cdot Red^{1.2}) \cdot (1 + 20 \cdot Green^{6.7})} - 0.32 \cdot f$$

$$\frac{\partial g}{\partial t} = 0.045 \cdot \frac{1}{(1 + 44 \cdot Red^{8.5}) \cdot (1 + 11 \cdot g^{7.8})} - 0.3 \cdot g$$

$$\frac{\partial h}{\partial t} = 0.1 \cdot \frac{(38 \cdot b^{8.6} \cdot 66 \cdot e^{8.6} + 38 \cdot b^{8.6} + 66 \cdot e^{8.6})}{(1 + 38 \cdot b^{8.6}) \cdot (1 + 66 \cdot e^{8.6}) \cdot (1 + 11 \cdot Red^{7.8}) \cdot (1 + 6.6 \cdot i^{1.8})} - 0.5 \cdot h$$

$$\frac{\partial i}{\partial t} = 0.06 \cdot \frac{\left((76 \cdot f^{1.3} \cdot 82 \cdot h^{8} + 76 \cdot f^{1.3} + 82 \cdot h^{8}) \cdot 25 \cdot i^{9.8} + (76 \cdot f^{1.3} \cdot 82 \cdot h^{8} + 76 \cdot f^{1.3} + 82 \cdot h^{8}) + 25 \cdot i^{9.8}\right) \cdot 8.4 \cdot Red^{9.3} \cdot 41 \cdot j^{5.9}}{(1 + 76 \cdot f^{1.3}) \cdot (1 + 82 \cdot h^{8}) \cdot (1 + 25 \cdot i^{9.8}) \cdot (1 + 8.4 \cdot Red^{9.3}) \cdot (1 + 41 \cdot j^{5.9})} - 0.16 \cdot i$$

$$\frac{\partial j}{\partial t} = 0.042 \cdot \frac{90 \cdot Green^{9.5}}{(1 + 90 \cdot Green^{9.5}) \cdot (1 + 7.4 \cdot Red^{4.6}) \cdot (1 + 99 \cdot d^{1}) \cdot (1 + 8.8 \cdot e^{1.1}) \cdot (1 + 88 \cdot g^{1.1})} - 0.12 \cdot j$$

$$\frac{\partial k}{\partial t} = 0.05 \cdot \frac{1}{(1 + 8.1 \cdot f^{1.4})} - 0.44 \cdot k$$

### Letter ‘C’ pattern (Fig. 8B):

$$\frac{\partial Blue}{\partial t} = 0.099 \cdot \frac{13 \cdot c^{9.8}}{(1 + 13 \cdot c^{9.8})} - 0.12 \cdot Blue$$

$$\frac{\partial a}{\partial t} = 0.0097 \cdot \frac{41 \cdot Red^{2.1} \cdot 45 \cdot Green^{9.4}}{(1 + 41 \cdot Red^{2.1}) \cdot (1 + 45 \cdot Green^{9.4}) \cdot (1 + 44 \cdot d^{3.5})} - 0.23 \cdot a$$

$$\frac{\partial b}{\partial t} = 0.093 \cdot \frac{24 \cdot Red^{3.7} \cdot 40 \cdot Green^{5.2} \cdot 61 \cdot e^{2.8}}{(1 + 24 \cdot Red^{3.7}) \cdot (1 + 40 \cdot Green^{5.2}) \cdot (1 + 76 \cdot d^{5.4}) \cdot (1 + 61 \cdot e^{2.8})} - 0.52 \cdot b$$

$$\frac{\partial c}{\partial t} = 0.093 \cdot \frac{48 \cdot a^{9.3}}{(1 + 3.2 \cdot Red^{3.5}) \cdot (1 + 48 \cdot a^{9.3}) \cdot (1 + 22 \cdot b^{9.4}) \cdot (1 + 60 \cdot d^{9.5})} - 0.27 \cdot c$$

$$\frac{\partial d}{\partial t} = 0.015 \cdot \frac{8.4 \cdot Green^{1.4}}{(1 + 8.4 \cdot Green^{1.4}) \cdot (1 + 3.2 \cdot b^{1.6})} - 0.69 \cdot d$$

$$\frac{\partial e}{\partial t} = 0.079 \cdot \frac{40 \cdot Green^{5.8}}{(1 + 40 \cdot Green^{5.8}) \cdot (1 + 7.6 \cdot Red^{2.5})} - 0.63 \cdot e$$
